# Virus-Host Interaction Gets *Lousy*: P1vir Phage Development Upon Impaired RNA Global Regulation in *Escherichia coli* Δ*hfq* Mutant Cells

**DOI:** 10.64898/2025.12.28.696737

**Authors:** Grzegorz Cech, Agnieszka Szalewska-Pałasz, Jakub Giełdon, Grzegorz Węgrzyn, Anna Kloska

## Abstract

Bacteriophage P1 is a classic tool in molecular genetics, and its virulent derivative, P1vir, is widely used for generalized transduction in *Escherichia coli*. Phage development depends on host physiology and metabolism, much of which is controlled by RNA-based regulation. The bacterial RNA chaperone Hfq is a global post-transcriptional regulator of stress and metabolic pathways, but its role in phage biology is poorly understood. Here, we examined the effect of Hfq loss on P1vir development by comparing the infection kinetics, virion morphology, and global transcriptomes of P1vir and its *E. coli* hosts in the wild-type and Δ*hfq* backgrounds. Deletion of *hfq* impaired P1vir lytic development, yielding smaller plaques, reduced burst size, and virions with smaller heads and thinner tails. P1vir transcriptional profiling showed that Δ*hfq*-specific dysregulation emerged early and intensified over time, with disrupted developmental timing, compromised replication and genome processing, and an unbalanced morphogenetic program. In wild-type cells, P1vir infection triggered broad, time-dependent reprogramming of host gene expression, including the induction of central carbon and amino acid metabolism and other anabolic pathways, consistent with a state that supports productive phage replication. In contrast, the Δ*hfq* mutant mounted a narrower response, with limited metabolic induction and late upregulation of chaperones and proteostasis factors, suggesting the accumulation of misfolded proteins or stalled assembly intermediates. Motif scanning of P1vir transcripts identified candidate Hfq-favored sequence features in several under-induced phage genes, suggesting direct Hfq–RNA regulation. Thus, Hfq, although not essential, is important for efficient P1vir lytic development by coordinating phage gene expression with host reprogramming.

## 1. Introduction

Bacteriophage P1vir, a virulent derivative of the temperate P1 phage that infects *Escherichia coli*, was first isolated nearly 60 years ago (1). P1vir carries a mutation in the *c*4−*ant1/2* region of its genome, leading to the constitutive production of the phage antirepressor protein (Ant) (2). As a result, the phage is unable to lysogenize and exclusively undergoes lytic development. P1 is highly effective for general transduction and is a valuable genetic tool for constructing bacterial strains (3,4). Moreover, given its broad host range and emerging use as a CRISPR–Cas9 delivery vehicle (5), further studies of bacteriophage P1 and its hosts are warranted to better exploit this system.

The development of P1vir, like that of other phages, strongly depends on the host cell machinery and is modulated by the host physiological state (6). In bacteria, metabolic and physiological adaptations to various stimuli are largely mediated by RNA (7). Hfq is an RNA chaperone that is a key factor in RNA metabolism and is considered a global regulator of gene expression in bacteria (8–10). Hfq has RNA-binding properties and modulates mRNA stability and translation (8). It also facilitates interactions between mRNA and small non-coding RNAs (ncRNAs), thereby regulating post-transcriptional gene expression (9). Hfq also interacts with other proteins, such as ribosomal proteins, RNA polymerase, and poly(A)polymerase (PAP I) (11,12). Direct Hfq-DNA interactions and Hfq-dependent control of DNA topology, replication, and transposition have also been described (13,14).

Although Hfq was initially identified as a host factor necessary for Qβ bacteriophage replication (15,16), the functional link between bacterial Hfq and bacteriophage development remains unexplored. It has been observed that lambda bacteriophages λ*c*I*857* and λvir form smaller plaques on *hfq* mutant lawns (17); however this phenomenon has not been fully discussed. Another study demonstrated that Hfq is involved in the rapid degradation of mRNA following infection with the phage T4 *dmt* mutant (18). Recently, several antisense sRNAs interacting with Hfq have been identified in bacteriophage-derived genomic regions of pathogenic *E. coli* strains (19) and phage-derived small RNAs have been proposed as additional layers of phage–host regulation (7,20). However, data on the role of Hfq in P1 phage development, especially considering the pleiotropic effects of Hfq on DNA and RNA metabolism, remain limited.

In this study, we examined for the first time how the loss of Hfq protein affects P1vir development in *E. coli* by comparing the infection kinetics in Δ*hfq* and wild-type (WT) strains and profiling the global transcriptomes of both hosts and the phage throughout the infection. By defining the Hfq-dependent changes in P1vir and host gene expression, we aimed to clarify how Hfq contributes to phage-driven reprogramming of the host during the lytic cycle and why P1vir development is impaired in the Δ*hfq* mutant.

## 2. Materials and methods

### 2.1. Bacterial strains, bacteriophage, plasmid

*E. coli* CF7968 (MG1655 but *rph^+^ ΔlacIZ, mluI*) strain, referred to as the wild type (21), was used in this work. The Δ*hfq* mutant strain (isogenic to CF7968) was used in this study (22). Bacterial strains were cultured at 37°C with shaking in Lysogeny Broth (LB) (Sigma-Aldrich) supplemented with 50 µg/ml kanamycin (Sigma-Aldrich) when required. The P1vir bacteriophage (1) and mini-P1 plasmid (pSP102; P1 replication *ori*; 1–2 copies/cell) (23) were used in this study.

### 2.2. Plaque morphology analysis

The plaque morphology of the P1vir bacteriophage was analyzed in wild-type and Δ*hfq* mutant cells. Serial dilutions of the P1vir bacteriophage lysates were prepared in TM buffer (10 mM Tris-HCl, pH 7.4, and 10 mM MgCl_2_). Next, 0.2 mL of the overnight culture was mixed with 0.3 mL of 100 mM CaCl_2_ in a 15 ml glass tube, and diluted P1vir lysate was added to obtain a final number of 10^2^–10^3^ phages per tube. Next, 3 mL of melted top agar (1% peptone, 0.5% NaCl, and 0.7% bacteriological agar) was added, mixed quickly, and poured onto freshly prepared LB agar plates.

The plates were then incubated overnight at 37°C. Plaque morphology was analyzed visually and using a binocular microscope. Plaque area and perimeter were measured using the Photoshop SC3 image analysis tool.

### 2.3. One-step growth experiment

Bacteria were cultured in LB to an optical density at 600 nm (OD_600_) of 0.2. Next, 10 ml of the culture was centrifuged at 4,000 × g. The pellet was resuspended in 1 ml of LB medium supplemented with 5 mM NaN_3_ and incubated for 5 min with shaking at 37°C. Next, the P1vir bacteriophage was added to a final multiplicity of infection (MOI) of 0.5, and the infected culture was immediately diluted 10,000 fold. Samples (0.5 ml) were withdrawn at the indicated time points, centrifuged, and the supernatant was titrated as described previously. Plaques were counted, and the phage yield and burst size were calculated.

### 2.4. Virion morphology analysis

High-titer P1vir lysates were prepared from 20 ml of culture. Polyethylene glycol (PEG) 8000 was added to the clarified lysates to a final concentration of 10%, and the samples were mixed at 4°C overnight. Precipitated phage particles were pelleted by centrifugation (8,000–10,000 rpm for 20–30 min) and resuspended in 15 ml of TM buffer (10 mM Tris-HCl, pH 7.4, 10 mM MgCl₂). A small amount of DNase was added before the chloroform extraction to remove residual DNA. Chloroform was then added, and the suspension was vigorously mixed and centrifuged to collect the aqueous phase. Chloroform extraction was repeated until the interphase “white layer” disappeared.

Purified phage lysates were examined using transmission electron microscopy (TEM) after negative staining. Briefly, a drop of lysate was applied to a carbon-coated TEM grid, allowed to adsorb for 3 min, and the excess liquid was removed. The grid was then stained with freshly prepared 1.5–2% (w/v) uranyl acetate for 30 s, blotted, and viewed under a Philips CM 100 transmission electron microscope. Virion dimensions were measured from the micrographs using Adobe Photoshop CS3 Extended.

### 2.5. Infection experiment and RNA isolation

Wild-type and Δ*hfq* strains were cultured in LB medium at 37°C with shaking to an OD_600_ of 0.2. Next, bacteriophage P1vir was added to the final MOI of 10. Samples (0.5 mL) were withdrawn 10 and 20 min after phage infection, mixed with RNAprotect^®^ Bacteria Reagent (Qiagen), according to the manufacturer’s protocol, and incubated for 5 min at room temperature (RT). A sample of bacterial culture before infection (denoted as time point 0) was withdrawn and processed similarly. RNA was extracted using the RNeasy^®^ Mini Kit (Qiagen) according to the manufacturer’s protocol, with an additional step of on-column DNase digestion using the RNase-Free DNase Set (Qiagen) to eliminate genomic DNA contamination. RNA was eluted with 50 µL of RNase-free water, and the column was washed twice with the same eluate to enrich the RNA yield. The purity and integrity of the RNA were assessed using a Bioanalyzer 2100 (Agilent Technologies, USA) and Agilent RNA 6000 Nano Kit (Agilent Technologies, USA). RNA samples were stored at −80°C prior to RNA sequencing.

### 2.6. Next-generation RNA sequencing and bioinformatic analysis

RNA sequencing (RNA-seq) was performed using the Illumina platform. Library preparation, bacterial ribosome reduction, and rapid run flow cell with paired-end 50-bp reads were performed at the Genomics Core Facility of the University of Alabama at Birmingham (UAB), Heflin Center for Human Genetics (Birmingham, AL, USA). *E. coli* K12 MG1655 genome (NCBI RefSeq NC_000913.3) and P1 phage genome (NCBI RefSeq NC_005856.1) were used as a reference genomes. Quality control was performed using the Trimmomatic software (24). Read sequences were filtered out with an average quality score of 20, which was required on a sliding window of five bases. Reads with lengths of less than 50 bp were rejected*. E. coli* rRNA sequences were filtered, and reads were mapped to the genome using Bowtie 2 software (25). PCR duplicates within the mapped reads were estimated using the Picard tools (26). PCR duplicates were not removed from the analysis. FeatureCounts software from the Subread package was used to count the paired-end reads that were uniquely mapped to individual genes (27). Differential expression analysis was performed using DESeq2 software (28) and summarized as log₂-transformed fold change (log2FC) in expression.

### 2.7. Functional gene annotation analysis

P1 gene functions were described based on the genome annotation by Łobocka et al. (6) with updates (29–31). For this study, phage genes were classified into the following functional modules: Regulation/Timing (early/late program coordination factors, transcriptional control, e.g., repressors/antirepressors, late promoter activators, regulatory RNAs, division-control genes), Replication/Processing (functions needed to replicate and process phage genome; replication, recombination, repair, DNA methylation, SSB/helicase-like factors), Packaging (DNA encapsidation machinery: terminase/packase subunits, portal protein, head maturation control), Morphogenesis (structural and assembly proteins for head, baseplate, tail, and fibers), Lysis (host cell exit cassette: holins, antiholins, endolysin, spanins), Maintenance/TA (elements that support plasmid-like maintenance or addiction systems), Immunity (superinfection, immunity systems), Other/Unknown (hypothetical proteins, IS-associated, tRNAs, lipoproteins) (full list of genes with module annotations in Supplementary **Table S1**).

Functional annotation analysis was performed using GSAn (32) for Gene Ontology (GO) to produce a concise, easy-to-discuss list of enriched GO terms, and DAVID (33,34) was used for KEGG pathways to provide pathway-centric information that complemented the GO results. Venn diagrams were constructed using the InteractiVenn online tool (35).

### 2.8. Measurement of mini-P1 plasmid DNA replication

Overnight bacterial cultures carrying the pSP102 plasmid were diluted 1:100 in fresh LB and incubated at 37°C. At defined time points, 1 ml samples were mixed with 10 μl [³H]-methyl-thymidine (1 mCi/ml; Hartmann Analytic) and incubated with shaking for 4 min at 37°C. Samples were then chilled on ice, centrifuged (10 min, 4000 × g, 4°C), and plasmid DNA was isolated using the GenElute Plasmid Miniprep Kit (Sigma-Aldrich). For each sample, 100 μl of the isolate was mixed with 2 ml scintillation fluid (PerkinElmer), and radioactivity was measured using a MicroBeta2 scintillation counter (PerkinElmer) and expressed as counts per minute (CPM).

### 2.9. Hfq-motif scanning

We extracted 5′ windows (−30..+60 nt) and 3′ windows (−40..+20 nt) around annotated coding sequences (CDSs, oriented to mRNA sense) from the P1 genome (NCBI GBFF). Motif definitions followed structural and biochemical studies that mapped Hfq-RNA interactions at the distal (A-rich AAN/ARN triplets), proximal 3’ U-rich tails), and rim (UA-rich patches) faces of Hfq (36–43). We scanned 5′ windows for (AAN)_n_ and (ARN)_n_ (n ≥ 3) and UA-rich repeats, and 3′ windows for poly U tracts (≥ 5 U), with these thresholds chosen in line with previous works linking clustered (AAN)_n_ and extended U-rich segments to high-affinity Hfq binding (36,40,42). The analysis was performed using Biopython 1.86 (44) (**Script 1**).

### 2.10. Statistics

The Shapiro–Wilk test was used to verify the normal distribution. Student’s *t*-test or the Mann–Whitney U test was used for two-sample comparisons using. Statistical significance was set at *p* < 0.05. The single-parameter bootstrap confidence interval for the population median was calculated using Stats.Blue online software (45) with a confidence level of 95%, and the number of bootstrap samples for the calculations was set to 10,000. Comparison of Hfq motif-positive and motif-negative transcripts was implemented using Biopython 1.86 (44) (**Script 2**).

## 3. Results

### 3.1. Impaired P1vir lytic development in wild-type and Δ*hfq* hosts

To assess how the loss of Hfq affects P1vir lytic development, we compared the plaque morphology, one-step growth kinetics, and burst size of phages released from wild-type and Δ*hfq E. coli* hosts.

P1vir formed visibly smaller plaques on Δ*hfq* cells than on wild-type cells (**Figure 1A**). The average plaque area was reduced by half (**Figure 1B**) and the perimeter by one-third (**Figure 1C**) compared to plaques formed on the wild-type strain. Additionally, the phage yield and burst size in Δ*hfq* cells were significantly lower than those in wild-type cells (**Figure 1D** and **1E**).

**Figure 1.**
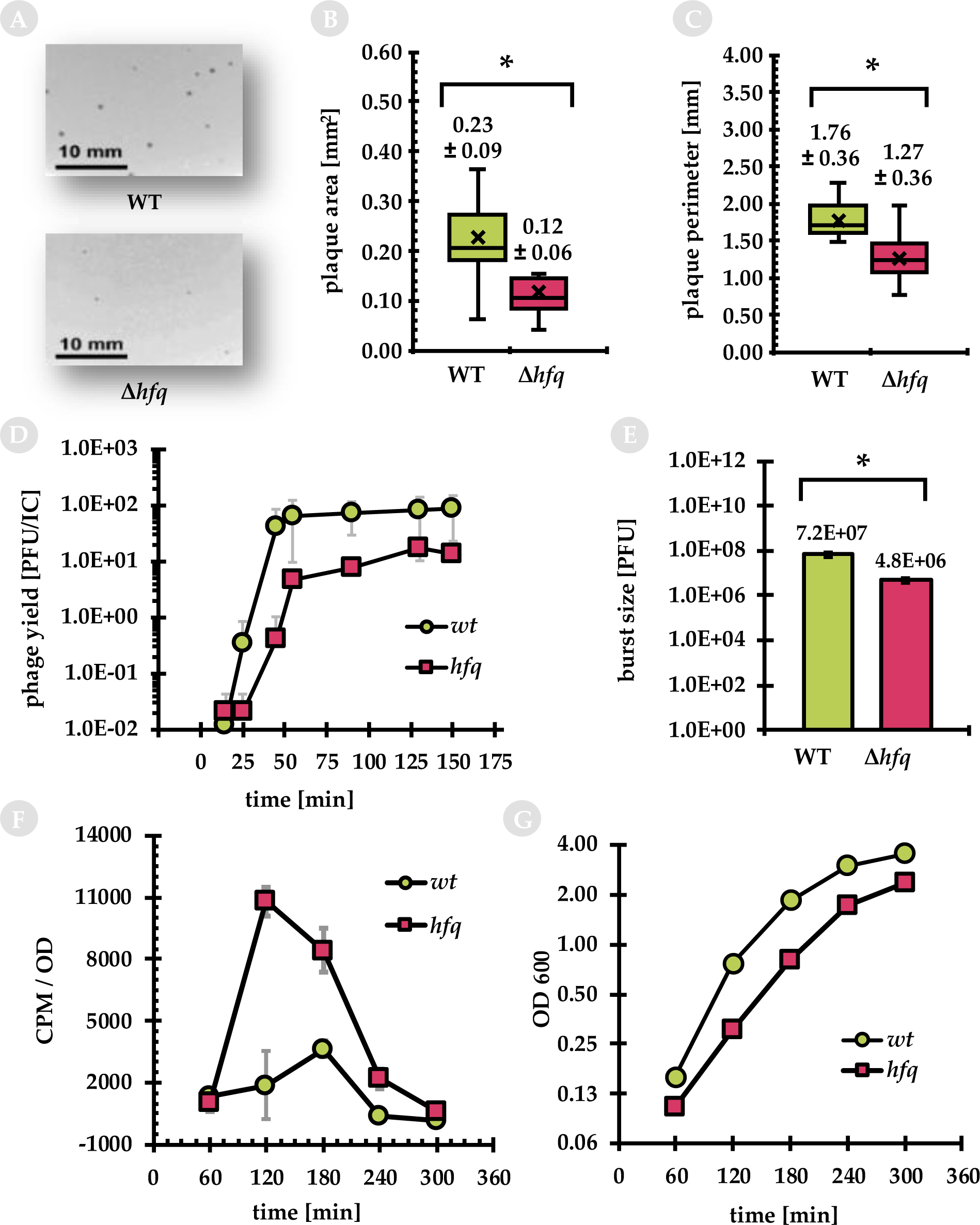
Development of P1*vir* phage in wild-type and Δ*hfq* host cells. (A) Plaque morphology of P1vir on bacterial lawn. (B) Area and (C) Perimeter of plaques formed by P1vir on the two hosts. Boxplots show the median (horizontal line), mean (×), interquartile range (box, Q1–Q3), and minimum–maximum (whiskers). Values are expressed as the mean ± SD. (D) Kinetics of P1vir development in a one-step growth experiment. Phage yield is reported as plaque-forming units (PFU) per infection center (IC) and is expressed as the mean ± SD from at least two independent experiments. (E) Burst size of phages released from bacterial hosts; results are presented as mean ± SD from four independent experiments. (F) Kinetics of P1 mini-P1 plasmid (pSP102) replication. Results of ^3^H-thymidine incorporation per density of bacterial culture. The results are presented as mean ± SD from three independent experiments. (G) Growth curves of bacteria harboring the mini-P1 plasmid. Statistics: * *p* < 0.05 (Student’s *t*-test). Abbreviations: CPM, counts per minute; IC, infection center; OD, optical density; PFU, plaque-forming unit; SD, standard deviation; WT, wild type.

### 3.2. P1vir virion morphology in wild-type and Δ*hfq* hosts

To assess whether the loss of Hfq affects virion assembly, we examined P1vir particles produced during lytic growth in wild-type and Δ*hfq* cells. High-titer lysates from both hosts were purified and analyzed using TEM, and individual virions were measured (head diameter, tail length, and tail width).

Virions produced in the Δ*hfq* mutant showed altered morphology and reduced size compared to those from the wild-type host; capsid head diameter and tail length and width were all significantly smaller (**Figure 2**). In lysates from both strains, we observed particles with contracted sheaths and visible central tail tubes (cores) for injecting DNA into a host cell, indicating that sheath contraction was preserved in the Δ*hfq* background. However, contracted particles appeared more frequently in the lysates from Δ*hfq*. Together, these observations suggest that P1vir morphogenesis is abnormal in the Δ*hfq* host cells.

**Figure 2.**
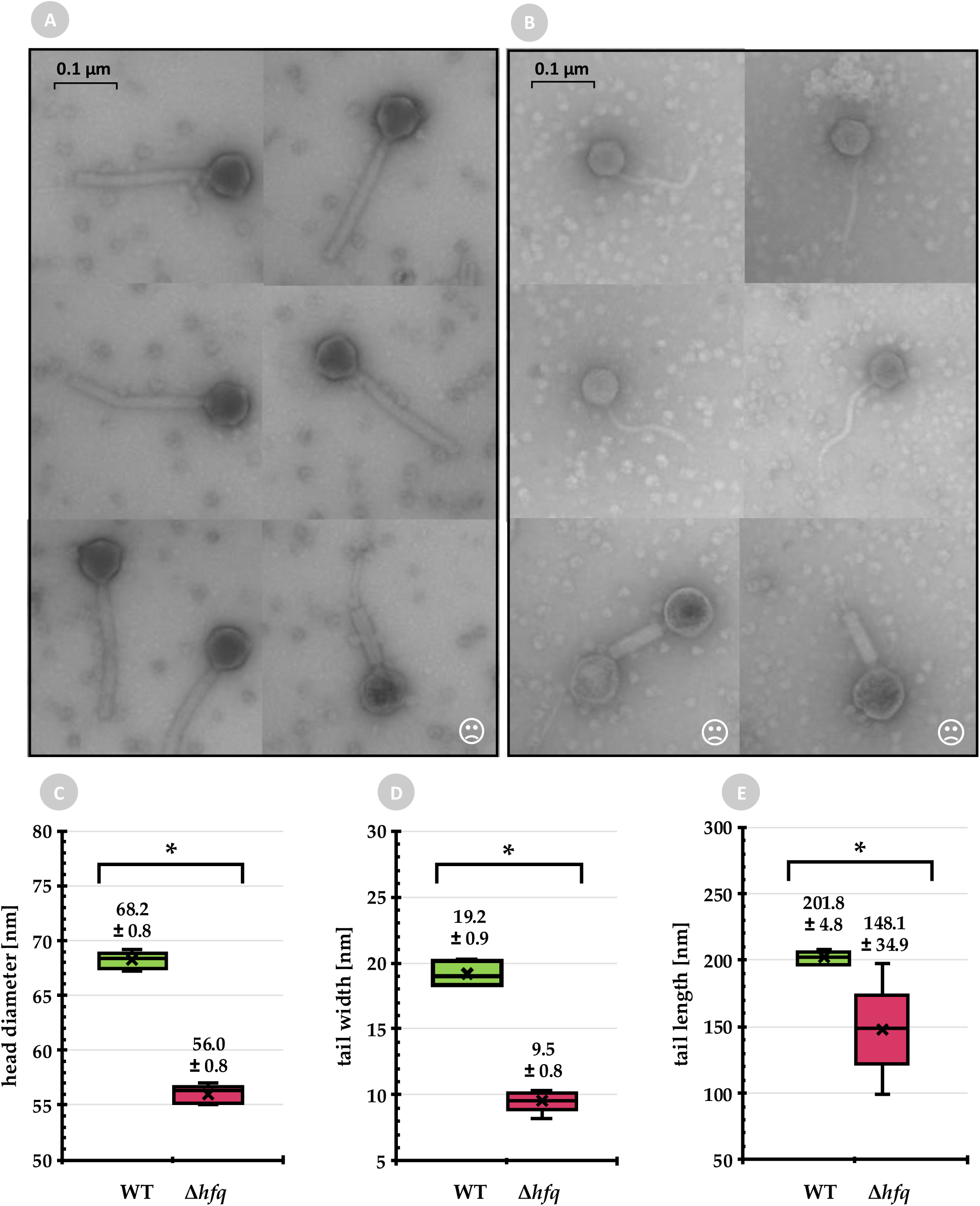
Morphology of P1vir virions produced in wild-type and Δ*hfq* host cells. Transmission electron microphotographs of virions produced in (A) the wild-type strain and (B) Δ*hfq* mutant. Quantification of (C) head diameter, (D) tail width, and (E) tail length of P1vir virions propagated in wild-type (green) and Δ*hfq* (red) cells. Boxplots show the median (horizontal line), mean (×), interquartile range (box, Q1–Q3), and minimum/maximum values (whiskers). The results are summarized as mean ± SD (values above the boxplots). Statistics: * *p* < 0.05 (Student’s *t*-test). Abbreviation: WT, wild type.

### 3.3. Global P1vir transcriptional program in wild-type and Δ*hfq* hosts

To understand which stages of the P1vir cycle are most affected by Hfq, we analyzed the global phage transcriptional program during infection in wild-type and Δ*hfq* cells.

We compared the phage transcriptomes of infections of wild-type and Δ*hfq* hosts. During P1 lytic development, some early phage genes are expressed as soon as 5 min post-infection, most early phage genes are induced between 10 and 15 min, and the expression of late genes begins between 20 and 30 min (6). Accordingly, we assessed gene expression at 10 min (early stage) and 20 min (late stage) post-infection using RNA-seq. Preliminary analysis showed no significant differences in phage gene expression between samples without phages (no phage) and those at 0 min after phage addition (Supplementary **Table S1**). Thus, for each phage gene, the log2 fold change (log2FC) in expression was calculated relative to the no-phage sample (indicated in this study as 0 min).

#### 3.3.1. Quantitative analysis of differentially expressed phage genes

Of the 117 phage genes, 38 and 45 were significantly (FDR < 0.05) differentially expressed (vs. 0 min) in the wild-type host at 10 and 20 min post-infection, respectively, while there were 39 and 43 genes in the Δ*hfq* host (**Figure 3A**). Genes that were significantly differentially expressed in at least one host or at one time point were included for functional analysis; thus, a total of 46 genes were included for further analysis.

**Figure 3.**
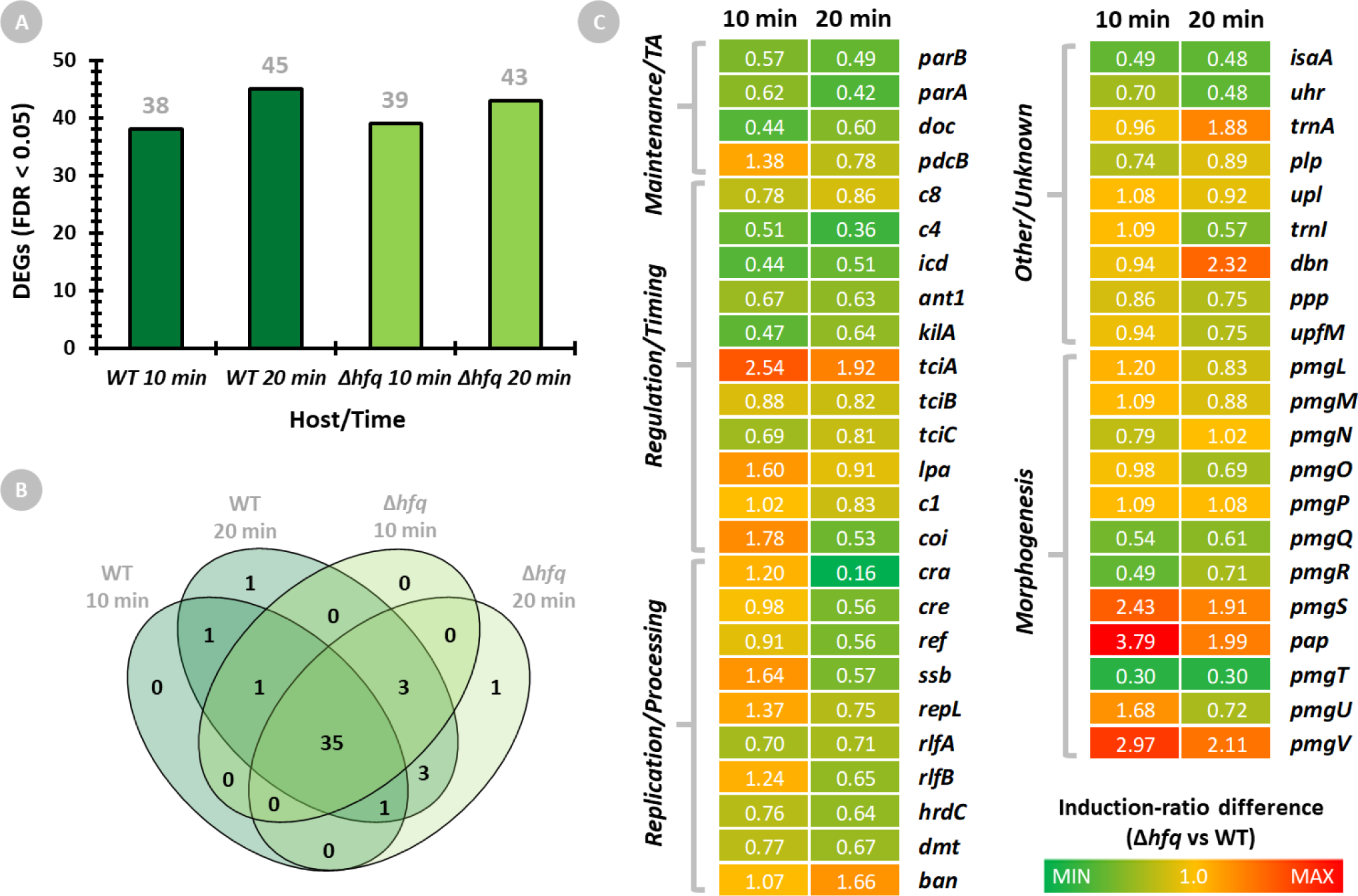
P1vir transcriptome during infection of wild-type and Δ*hfq* host cells. Phage transcriptomes isolated from wild-type and Δ*hfq* cells were profiled using RNA sequencing at 10 min and 20 min time points. Differential expression was summarized as log2 fold change (log2FC) relative to the 0 min time point. (A) Number of significantly upregulated phage genes (FDR < 0.05; vs. 0 min) at each time point in each host. (B) Venn diagram showing the overlap of upregulated phage genes (vs. 0 min) across hosts and time points. (C) Relative expression of P1*vir* genes during infection of the Δ*hfq* host vs. WT. The heat map shows the induction ratio difference (IRD) for 10 and 20 min, defined as IRD = 2^ΔΔlog2FC^, where ΔΔlog2FC = (log2FC_Δ*hfq*_ − log2FC_WT_) at the same time point Genes are annotated by functional module. Abbreviations: DEGs, differentially expressed genes; FDR, false discovery rate; TA, toxin–antitoxin; WT, wild type.

To compare phage gene expression between Δ*hfq* and wild-type hosts at 10 and 20 min post-infection, we computed a between-strain contrast at each time point (ΔΔlog2FC) according to Equation (1): ΔΔlog2FC*_t_* = (log2FC_hfq,*t* vs. 0_ − log2FC_WT,*t* vs. 0_). The size of induction-ratio difference (IRD) in phage gene expression in Δ*hfq* vs. wild-type was calculated as 2^ΔΔlog2FC^ to determine how much stronger (IRD > 1) or weaker (IRD < 1) the infection-induced change is in Δ*hfq* compared to wild-type at the specific time point.

In both hosts, P1vir expressed essentially the same set of genes at 10 and 20 min, with 43 genes overlapping between strains (**Figure 3B**, Supplementary **Table S2**), but the expression levels diverged markedly between strains (Supplementary **Table S1**). For example, using an IRD threshold of ≤ 0.5 or ≥ 1.5 (i.e., ≥ 50% lower or ≥ 50% higher gene induction in Δ*hfq* vs. WT), at 10 min, six genes were substantially less induced (IRD ≤ 0.5) and eight were more induced (IRD ≥ 1.5) in Δ*hfq*; at 20 min, seven genes were less induced and seven were more induced. With a more permissive window, the number of under- or over-induced phage genes in Δ*hfq* was even higher.

#### 3.3.2. Qualitative analysis of differentially expressed phage genes

For functional analysis of the phage transcriptome, phage genes were first assigned to functional modules: Regulation/Timing, Replication/Processing, Packaging, Morphogenesis, Lysis, Maintenance/TA, Immunity, and Other/Unknown. To characterize the biological pattern of phage transcriptome changes, we estimated the module-level effect size summary (Δ*hfq* relative to WT) by calculating the per-module medians of ΔΔlog2FC (with 95% CI) at each time point (**Table 1**).

**Table 1.** Module-level effect size summary of P1vir gene expression differences between Δ*hfq* and wild-type hosts. For each functional module, the median ΔΔlog2FC (95% CI) at 10 and 20 min post-infection across significantly differentially expressed genes in that module was calculated. Negative module medians indicate under-induction, whereas positive medians indicate over-induction in Δ*hfq* relative to WT. *N* reports the number of significantly differentially expressed (vs. 0 min) phage genes annotated to each module. Abbreviations: CI, confidence interval; FC, fold change; *N*, number of genes; TA, toxin–antitoxin; WT, wild type.

| Module | *N* | Median $\Delta\Delta\log_2FC$ (95% CI)<sup>1</sup> | |
| --- | --- | --- | --- |
|  |  | 10 min | 20 min |
| Maintenance/TA | 4 | −0.751 (−1.1879, 0.4675) | −0.889 (−1.2353, −0.3652) |
| Regulation/Timing | 11 | −0.350 (−0.9762, 0.6791) | −0.303 (−0.9291, −0.2104) |
| Replication/Processing | 10 | 0.039 (−0.3696, 0.3056) | −0.632 (−0.8246, −0.4943) |
| Other/Unknown | 9 | −0.088 (−0.5148, 0.1073) | −0.417 (−1.0711, 0.9096) |
| Morphogenesis | 12 | 0.124 (−0.6112, 1.0153) | −0.226 (−0.5145, 0.5229) |

<sup>1</sup> Median $\Delta\Delta\log_2FC$ was calculated per module across significantly differentially expressed phage genes at the same time point, using single-gene values defined as $\Delta\Delta\log_2FC = (\log_2FC_{\Delta hfq} - \log_2FC_{WT})$ , with each $\log_2FC$ calculated relative to 0 min); CI was estimated by bootstrap percentile (10,000 resamples).

The largest negative module medians (ΔΔlog2FC; Δ*hfq* vs WT) among significant genes revealed clear transcript deficits in Maintenance/TA, Regulation/Timing, and Replication/Processing when P1vir developed in Δ*hfq*. Within these functional modules, the expression levels of phage genes differed markedly between strains and across time points, revealing notable differences in the developmental patterns between the Δ*hfq* and wild-type hosts (**Figure 3C**).

In Maintenance/TA, components involved in plasmid partitioning (*parA*/*parB*) and toxin–antitoxin functions (*doc*) were consistently under-induced in Δ*hfq* cells. Interestingly, *pdcB* (post-doc locus) was transiently elevated at 10 min and attenuated at 20 min. Together, these results suggest weakened or impaired stabilization of the P1 genome during the lytic program.

In terms of Regulation/Timing, Δ*hfq* exhibited clear mis-timing of the expression of a broad range of factors involved in early/late program coordination and transcriptional control. At 10 min, *lpa* and *coi* showed transient boosts, in contrast to the normal level of *c1* (master repressor maintaining lysogeny). Small regulatory RNA *c4,* which modulates immunity and late-gene circuitry, shows persistent under-induction. By 20 min, most regulators, including *c1*, are reduced relative to the wild type, indicating a weakened late regulatory program and a disrupted early → late switch in the absence of Hfq. Moreover, cell division-control genes also showed deregulation in the Δ*hfq* host, with *tciA* over-induced and *icd*, *ant1*, *kilA*, and *tciB*/*C* under-induced at both time points, suggesting that the coordination of host physiology with phage development may be mis-timed or deregulated/diminished.

Replication/Processing was slightly induced at 10 min (positive module median ΔΔlog2FC) with a few early transcription boosts in Δ*hfq* for early lytic replication factors (*repL*, *rlfB*) and DNA-binding protein (*ssb*). Interestingly, the putative recombination factor (*hrdC*) and N6-adenine DNA methyltransferase (*dmt*) were persistently under-induced. By 20 min, the module median ΔΔlog2FC shifted broadly negative, with almost all genes (*cra*, *cre*, *ref*, *ssb*, *repL*, *rlfA*/*B*, *hrdC*, and *dmt*) showing under-induction. *Ban* was the only overexpressed exception. These results suggest a transient boost in the activation of early replication, followed by a late-stage deficit in replication in Δ*hfq*. In addition, deficits in DNA methylation and recombination suggest disrupted genome processing, influencing further replication or packaging competence.

Morphogenesis showed mixed signals, with the module median ΔΔlog2FC being slightly elevated at 10 min but shifting modestly negative by 20 min in Δ*hfq*. Here, the Δ*hfq* host permits selective early over-induction of some morphogenetic genes (*pmgS*/*V*/*U*, *pap*), whereas many others are reduced (*pmgQ/R/T*). By 20 min, some early elevated genes remain over-expressed (*pmgS*/*V*, *pap*), whereas others become under-expressed (*pmgO*/*U*), yielding an unbalanced structural program, suggesting assembly discoordination, which is in concordance with altered P1vir virion morphology in Δ*hfq*.

The expression profile of the Other/Unknown category was mixed. For example, tRNA genes diverged (*trnA*↑, *trnI↓* at 20 min), and several uncharacterized loci were strongly deregulated (*uhr*↓, *dbn*↑ at 20 min) in the Δ*hfq* strain.

Together, these patterns indicate that the loss of Hfq substantially reprograms P1vir transcription, underscoring the key role of Hfq in productive phage development. At the transcriptome level, we observed deficits in replication and DNA processing, disrupted developmental timing, an unbalanced morphogenetic program, and weakened genome maintenance, providing a coherent transcriptional basis for the reduced burst size and small plaques of P1vir in the absence of Hfq.

### 3.4. Replication of mini-P1 plasmid in wild-type and Δ*hfq* hosts

Because our phage transcriptome data indicated deregulation of replication and maintenance genes in the Δ*hfq* host, we next examined the replication of a mini-P1 plasmid to test whether Hfq influences P1 origin-dependent DNA replication independently of the full lytic cycle. The mini-P1 plasmid is a P1-derived replicon that retains the phage origin and replication/partitioning functions but lacks lytic and structural genes; therefore, it is maintained as a low-copy P1-based plasmid in *E. coli*.

We assessed mini-P1 (pSP102) replication kinetics in wild-type and Δ*hfq* strains during growth from 60 to 300 min by measuring the incorporation of [^3^H]-thymidine. At the onset of exponential growth, the Δ*hfq* mutant showed a markedly higher mini-P1 replication efficiency than the wild-type strain (**Figure 1F–G**). As the cultures progressed, the replication efficiency declined in both strains. These results indicate that replication from the P1 origin is more efficient in the absence of Hfq.

### 3.5. Host transcriptional response to P1vir infection in wild-type and Δ*hfq* strains

Because phage development depends critically on host physiology, we examined how wild-type and Δ*hfq E. coli* cells transcriptionally respond to P1vir infection over time.

Host gene expression was measured using RNA-seq at 0 min (pre-infection), 10 min (early), and 20 min (late) post-infection. To focus on infection-induced changes within each genotype, expression values for each strain were normalized to the corresponding 0-min (pre-infection; OD_600_ = 0.20) sample, which removes baseline transcriptomic differences between strains and yields fold changes attributable to infection. We assumed that incubating non-infected control cultures for an additional 10–20 min would not produce major transcriptomic changes; therefore, the detected changes were attributed to P1vir infection.

#### 3.5.1. Quantitative analysis of differentially expressed host genes

Upon P1vir infection, the number of significantly (FDR < 0.05) differentially expressed host genes differed markedly between strains and time points (**Figure 4A**, Supplementary **Table S3**). In the wild-type strain, 115 and 400 genes were significantly differentially expressed at 10 and 20 min post-infection, respectively, whereas in the Δ*hfq* mutant, only 34 and 45 genes were deregulated at these time points. The vast majority of genes that were upregulated or downregulated in response to infection were specific to the wild-type strain and were not differentially expressed in the Δ*hfq* mutant (**Figure 4B-C**, Supplementary **Table S4**). Notably, eight genes were exclusively downregulated and 13 were upregulated in the Δ*hfq* mutant upon infection.

**Figure 4.**
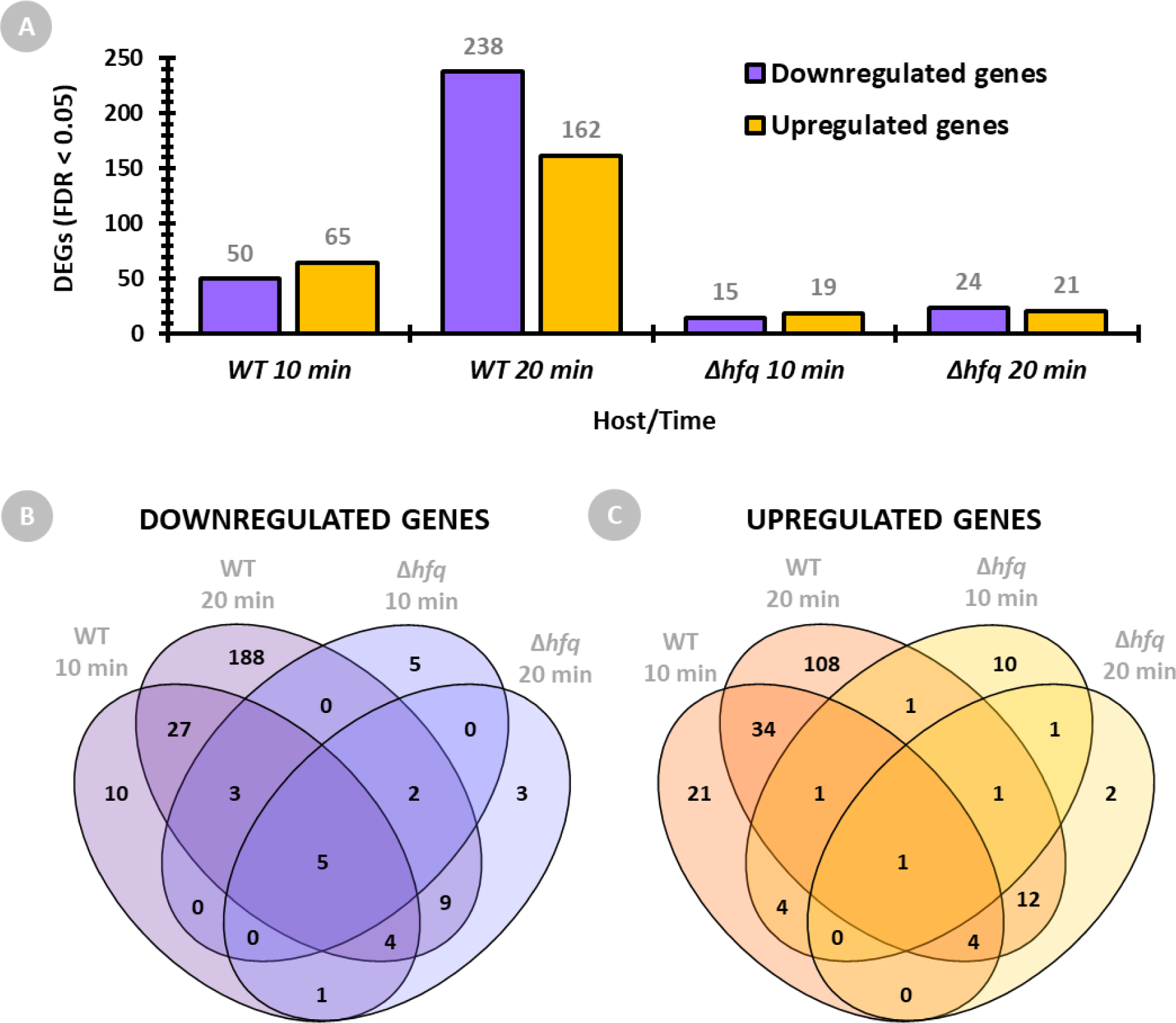
Global host gene expression changes following P1vir infection. Wild-type and Δ*hfq* host transcriptomes were analyzed using RNA sequencing. Differential gene expression was assessed at 10 and 20 min post-infection compared to that at 0 min. (A) Number of significantly (FDR < 0.05) downregulated and upregulated genes detected in hosts and time points. Venn diagrams showing the overlap of (B) downregulated and (C) upregulated genes between the hosts and time points. Abbreviations: DEGs, differentially expressed genes; FC, fold change; FDR, false discovery rate; WT, wild type.

#### 3.5.2. Qualitative analysis of differentially expressed host genes—top responders

The top differentially expressed host genes (**Figure 5**; Supplementary **Table S5**) revealed distinct host-dependent programs during P1vir infection. In wild-type cells, the response at 10 min was dominated by the reprogramming of metabolism and nutrient transport, notably the induction of glutamine uptake and metabolism (*glnH*/*A*/*P*/*Q*, *metN*) and sugar transport, followed by the engagement of phosphate (*pstC*/*S*) and stress/protein quality control pathways (e.g., *clpB*, *ibpA*) at 20 min. In contrast, the Δ*hfq* mutant showed early induction of sulfur amino acid metabolism, especially glutamine and methionine pathways (*glnH*, *glmS*, *and met* genes), followed by a pronounced upregulation of chaperones and proteostasis factors (*dnaK*, *groL*/*S*, *clpB*, *hslU*/*V*, and *grpE*) at 20 min.

**Figure 5.**
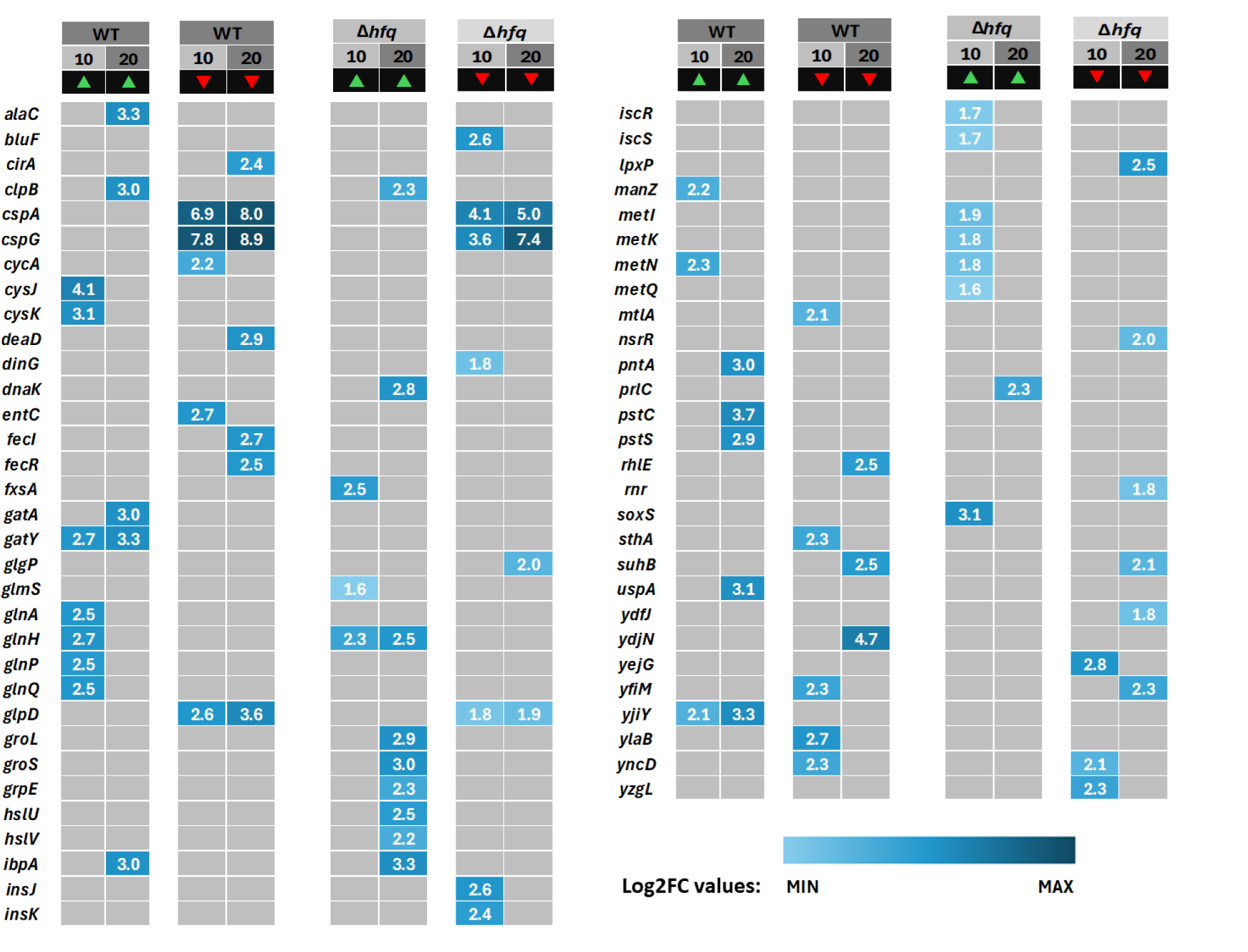
Top ten most differentially expressed host genes following P1vir infection. The heat map displays the ten most downregulated (red ▾) and ten most upregulated (green ▴) genes at 10 and 20 min post-infection for both Δ*hfq* and wild-type (WT) strains. Gene expression is reported as log2 fold change (log2FC) relative to that at 0 min. The gene names and ranks are listed in Supplementary **Table S5**.

Functionally, these differences indicate that the wild-type host mounts a broad, coordinated metabolic reprogramming to support or respond to infection, whereas loss of Hfq blunts this program and shifts the cell toward stress- and proteostasis-focused responses. Notably, across both strains and time points, *cspA*/*cspG* and *glpD* were consistently downregulated, indicating an infection-associated suppression of cold-shock programs and glycerol-3-phosphate oxidation, independent of Hfq status.

#### 3.5.3. Functional analysis of differentially expressed host genes

Functional enrichment analysis was performed on datasets of upregulated and downregulated host genes that were significantly differentially expressed (FDR < 0.05). Gene Ontology (GO) enrichment across Biological Process (BP), Cellular Component (CC), and Molecular Function (MF) categories was identified with GSAn tool. KEGG pathway enrichment was identified using the DAVID tool. For KEGG results, the fold-enrichment score was used to indicate term specificity. In the GSAn output, the information content (IC) score was used as an analogous measure of specificity.

Based on KEGG pathway analysis (**Figure 6**), P1vir infection in wild-type cells triggered broad metabolic rewiring, with seven pathways downregulated and 23 pathways upregulated; several overlapped across time points, with more genes affected at 20 min. In contrast, the Δ*hfq* mutant showed a much narrower response, with only two pathways downregulated and three upregulated, each involving fewer genes. Consistent with KEGG, GO enrichment (**Figure 7**) also showed fewer deregulated terms in Δ*hfq* than in wild-type at both time points—11 versus 22 terms at 10 min post-infection and 12 versus 43 at 20 min post-infection.

**Figure 6.**
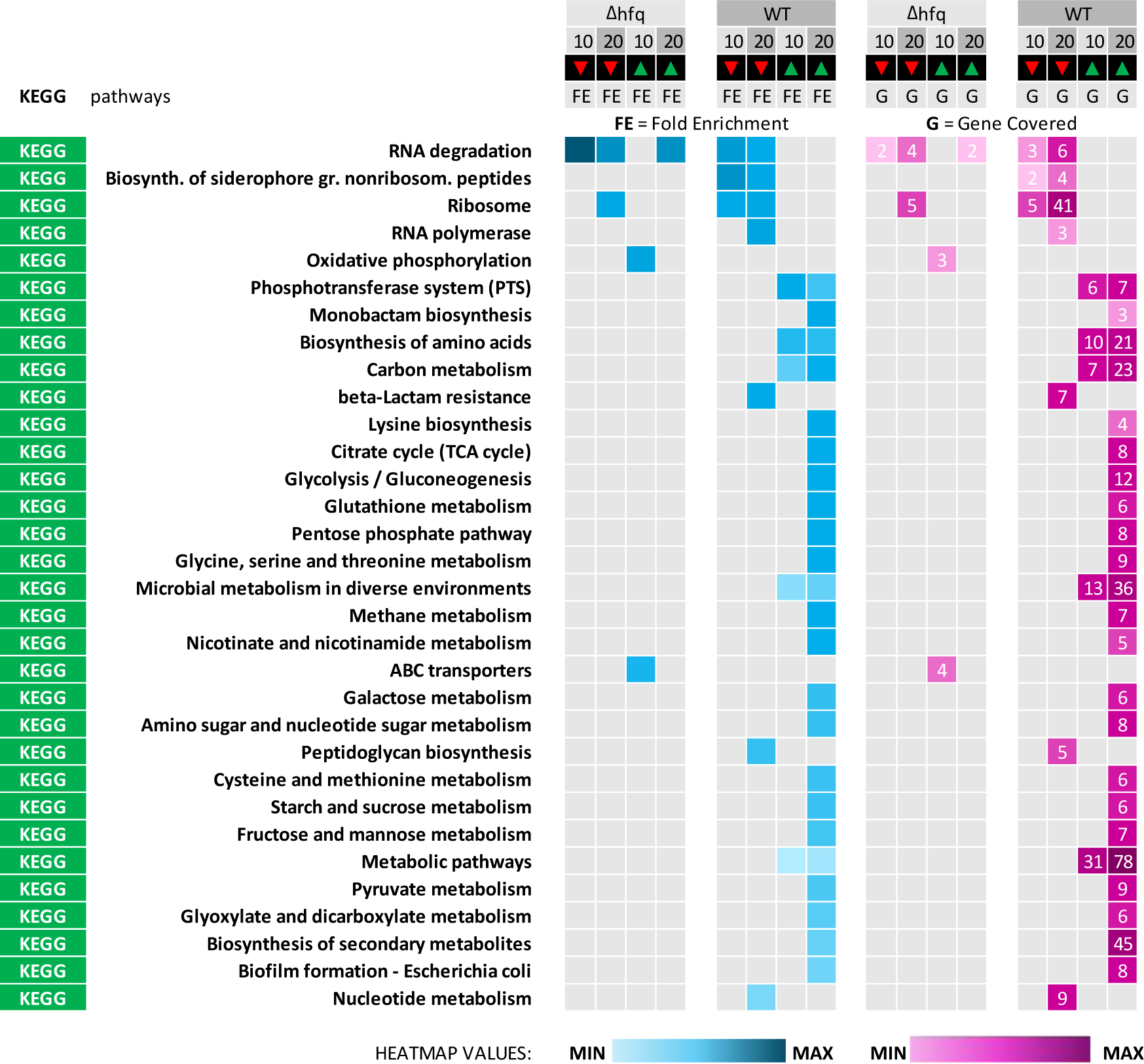
KEGG pathway analysis of host transcriptomes following P1vir infection. KEGG pathway enrichment was performed using DAVID for datasets of downregulated and upregulated genes (red ▾and green ▴, respectively) in wild-type (WT) and Δ*hfq* hosts at 10 and 20 min post-infection. Terms were ordered by the row-wise sum of fold-enrichment scores across hosts/time points (blue heatmap; higher scores indicate greater term specificity). The corresponding number of differentially expressed genes annotated to each term (pink heatmap; numbers represent the genes covered) is shown.

**Figure 7.**
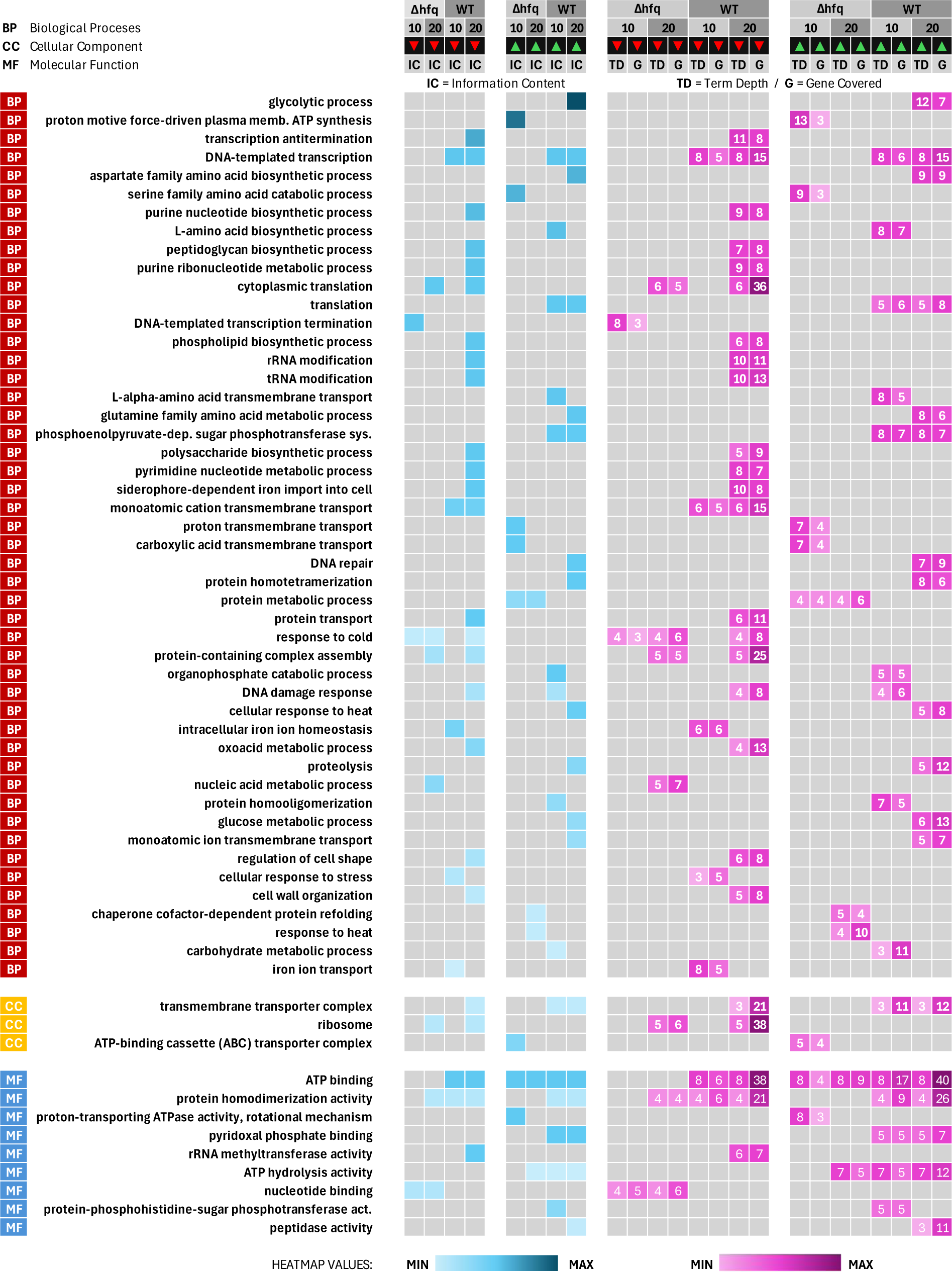
Gene Ontology (GO) term analysis of host transcriptomes following P1vir infection. GO enrichment was performed using the GSAn tool for datasets of down- and upregulated genes (red ▾and green ▴, respectively) in wild-type (WT) and Δ*hfq* hosts at 10 and 20 min post-infection. Specific GO terms from the three ontologies—Biological Process (BP), Molecular Function (MF), and Cellular Component (CC) are shown. Terms were ordered by the row-wise sum of Information Content (IC) scores across hosts/time points (blue heatmap; darker colors represent higher scores, i.e., greater specificity). Term depth (TD; pink heatmap; numbers represent the level of detail a GO term provides) and the corresponding number of annotated differentially expressed genes (G; pink heatmap; numbers represent the number of genes covered) are shown.

Both enrichment analyses showed that P1vir infection induced a broad, time-dependent response in the wild-type host, whereas the Δ*hfq* mutant elicited a much narrower program. In wild-type cells, KEGG enrichment (**Figure 6**, Supplementary **Table S6**) indicated coordinated upregulation of central metabolism at 10 min (phosphotransferase system (PTS), amino acid biosynthesis) that broadened by 20 min (tricarboxylic acid (TCA) cycle, glycolysis/gluconeogenesis, pentose-phosphate pathway) alongside sustained downregulation of ribosome and RNA degradation and additional repression of RNA polymerase, nucleotide metabolism, and peptidoglycan biosynthesis. This pattern suggests that the wild-type host redirects resources toward energy production and biosynthesis with concurrent remodeling of gene expression and cell wall pathways. In contrast, the Δ*hfq* host showed a narrow KEGG response, which included early enrichment of oxidative phosphorylation and ABC transporters, with only limited RNA degradation and ribosome-related changes detected.

The GO enrichment results (**Figure 7**, Supplementary **Table S7**) corroborate and refine these trends. At 10 min, the wild-type host showed coordinated induction of biosynthesis and transport (BP: amino acid biosynthesis and transport, sugar PTS; CC: transmembrane transporter complex) with ATP binding and hydrolysis functions, whereas selected iron homeostasis, transport, and transcription terms decreased. By 20 min, DNA repair and central carbon metabolism (glucose metabolic process, glycolysis, TCA, and glutamine-family amino acid metabolism) were induced, whereas nucleotide and peptidoglycan biosynthesis, cell wall organization, and ribosomal and translation terms were reduced. In Δ*hfq*, early upregulated GO terms centered on energy generation and membrane transport (e.g., BP: proton-motive-force-driven ATP synthesis, proton transmembrane transport, MF: ATP binding, and ATPase activity; CC: ABC transporter complex) together with amino acid metabolism (BP: serine-family amino-acid catabolism), while response to cold (BP) decreased. By 20 min, nucleic acid metabolism, translation, and ribosomes were downregulated (BP: nucleic acid metabolic process, cytoplasmic translation; CC: ribosome), and chaperone-dependent protein refolding (BP) and ATP hydrolysis activities were increased, indicating a late response dominated by proteostasis.

To sum up, the wild-type host exhibited a progressive increase in transcriptional responsiveness during the course of infection, whereas this response was severely impaired in the Δ*hfq* mutant. Overall, Δ*hfq* lacks the extensive metabolic induction observed in the wild-type strain following P1vir infection. Wild-type cells activate central carbon and amino acid metabolism while selectively suppressing parts of the translation and RNA turnover machinery. The Δ*hfq* host displayed an early energy and transport response, followed by a late-stage proteostasis shift.

### 3.6. Hfq-favored motif features in P1vir transcripts

To explore whether some of the Hfq-dependent transcriptional effects on P1vir reflect direct Hfq–RNA interactions, we scanned phage transcripts for sequence features characteristic of Hfq-favored binding motifs. RNA fragments were scanned for the following Hfq-sensitive motifs: repeated AAN/ARN triplets, poly U tract, and UA-rich patches. A transcript was motif-positive if it had ≥ 3 consecutive repeats of the motif (AAN/ARN/UA) or had a 3′ poly-U tract length ≥ 5 nt. To test whether these Hfq-favored features were associated with stronger transcriptional defects in Δ*hfq*, we compared ΔΔlog2FC (Δ*hfq* vs. WT) at 10 and 20 min of motif-positive transcripts to motif-negative transcripts using Mann–Whitney U (MWU) test.

Several strongly under-induced in Δ*hfq* phage transcripts carry extended AAN/ARN clusters or 3′ poly-U tracts around P1 start and stop codons, suggesting that they are potentially sensitive to Hfq-dependent regulation (**Table 2**; Supplementary **Table S8** for complete motif-scan dataset).

**Table 2.**
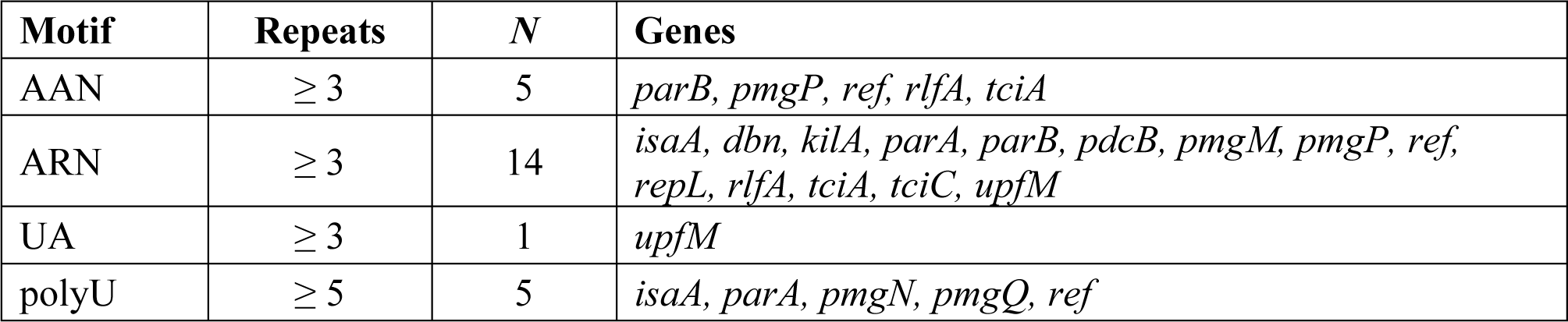
Candidate Hfq-favored P1vir transcripts. Phage genes were selected based on the presence of strong Hfq-favored sequence motifs: (AAN)_n_, (ARN)_n_, or (UA)_n_, where n ≥ 3 consecutive repeats in the 5′ window, and poly U tracts where U ≥ 5 nt in the 3′ window. The complete motif scan results are provided in Supplementary **Table S8**.

Transcripts with 3′ poly-U tracts (polyU ≥ 5 nt) showed a trend toward stronger under-induction (medians at 10 min of −0.70 vs. −0.03; MWU *p* = 0.059; at 20 min of −0.82 vs. −0.42; *p* = 0.101), but this did not reach statistical significance, likely reflecting the small number of P1 genes. The AAN, ARN, and UA clusters showed no detectable association with stronger under-induction (all *p* >> 0.1; Supplementary **Table S9**). Therefore, we treated these motif-based associations as hypothesis-generating rather than definitive evidence for direct Hfq–RNA regulation at specific P1vir loci.

## 4. Discussion

In this study, we show that the host Hfq protein is required for a properly timed and productive P1vir lytic program and for broad host reprogramming during infection. Deletion of *hfq* impairs P1vir development, yielding smaller plaques, reduced burst size, and altered virion morphology. Other phages, such as Qβ and λvir, are also sensitive to the loss of Hfq. Hfq is required for Qβ RNA replication (15,46) and efficient λvir development (λvir forms fewer and variable-sized plaques on *hfq* mutant cells) (17). Thus, our finding that P1vir development is similarly compromised in Δ*hfq E. coli* places P1vir in a broader class of phages whose lytic efficiency depends on Hfq. Global P1vir transcriptional profiling showed that Δ*hfq*-specific dysregulation emerged early (10 min) and intensified by 20 min, affecting developmental regulation and timing, replication, genome processing and maintenance, and morphogenesis. Moreover, phage–host interaction is impaired, resulting in blunted Δ*hfq* host reprogramming.

### P1vir program in the Δhfq mutant

The most prominent phage defect in the Δ*hfq* host concerns the regulation and timing of events during the lytic cycle. First, we have early over-induction of the late promoter activator locus (*lpa*), and Lpa activates late transcription once lytic replication is underway (47). However, we see a mistimed early pulse of its expression, which can perturb the onset of transcription from late promoters (e.g., morphogenesis and packaging genes), resulting in an improper early → late switch. We also have early over-induction and late under-induction of *coi* coding for the major C1 antirepressor (Coi) responsible for the lysis switch (48). Altered *coi*/*c1* levels may perturb promoter usage during the lytic cycle. The persistent downregulation of two other regulatory loci, *c4* and *ant1*, reflects a broader collapse of regulatory coordination. c4 is a small antisense regulatory RNA that pairs with ant mRNA and represses the synthesis of the Ant antirepressor (49). Normally, Ant binds to and inactivates the C1 repressor, thereby turning off C1-mediated repression of lytic genes. P1vir is characterized by the constitutive production of the antirepressor protein Ant due to mutations in the *c4*−*ant1*/*2* region (2,50). Ant continuously antagonizes C1 repression, thereby blocking lysogeny and restricting P1vir to the lytic cycle.

However, the Ant level still depends on transcription and RNA stability. The persistent under-induction of *c4* and *ant1* in Δ*hfq* suggests that even in the P1vir background, the c4–Ant regulatory module fails to reach its normal lytic output, weakening C1 antagonism and late promoter drive, thereby contributing to the globally dampened late program we observe. Finally, the misregulation of cell division control genes (*icd*, *kilA*, *tciA*/*B*/*C*) in Δ*hfq* cells points to impaired coordination of phage–host interactions, possibly by improper timing of host growth arrest, which would ordinarily support a productive phage program.

Early misregulation of core replication and processing genes suggests a brief, uncoordinated burst of replication in Δ*hfq* cells. Early over-induction of core replication genes, such as *repL* (initiator for lytic replication from *ori*L) and *ssb* (single-stranded DNA-binding protein), may be involved in the early boost of replication efficiency. By 20 min, these transcript levels drop more than in the wild-type host, but at this time, the *ban* gene coding for replicative DNA helicase (DnaB analog (51)) is over-induced, suggesting some mis-timing or compensatory mechanism to sustain late replication. Additionally, persistent under-induction of DNA processing and modification loci (*hrdC*—recombination, *dmt*—methylation (6)) is likely to further compromise replication efficiency.

Beyond its role in RNA metabolism, Hfq also affects DNA transactions. Hfq was shown to bind single-stranded DNA (ssDNA) via its amyloid-like C-terminal domain (CTR) and to alter ssDNA structure suggesting a direct role in managing ssDNA during DNA replication and recombination (52). Consistently, Hfq deficiency was previously shown to affect ColE1-like, but not λ-derived, plasmid replication (22). Here, the mini-P1 replication is more efficient in Δ*hfq* early in exponential growth, which might reflect the stronger early over-induction of key replication factors (*repL*, *rlfB*, *ssb*). However, in full P1vir infection, this boost is transient and is followed by a broad under-induction of many replication and processing genes, suggesting that productive replication requires tight coordination with recombination and methylation. Thus, we propose that the lack of Hfq may transiently accelerate early phage replication, but this burst is uncoordinated and is followed by the collapse of the replication program at a later stage. Such discoordination of genome replication and processing likely affects the pool of genomes available for packaging.

This picture of transcriptional deregulation is complemented by an unbalanced morphogenetic program, and misregulated *pmg*-family (putative morphogenetic function) transcript levels likely distort the stoichiometry of structural proteins. Previously published mutational analyses of the *pmg* cluster indicate that PmgA, PmgB, and PmgG are essential for tail morphogenesis, whereas PmgC and PmgR are required for head completion or head–tail attachment, as deletions in these genes produce aberrant or incomplete particles and fail to yield infectious virions (53). Here we hypothesize that this may be caused by an early *lpa* boost leading to premature and mismatched activation of the *pmg* cluster, so that some morphogenesis genes are expressed too early or in excess, while others are substantially reduced in expression. This imbalance is expected to cause assembly discoordination and ultimately inefficient virion production, consistent with the altered P1vir virion morphology observed by TEM (smaller heads and thinner tails and more frequent contracted sheaths in the Δ*hfq* host).

Taken together, we propose that in the complete P1vir lytic cycle, loss of Hfq leads to disrupted timing of developmental regulation, uncoordinated replication and genome processing, and unbalanced morphogenesis, collectively reducing infection efficiency and yielding the reduced burst size and smaller plaques observed in Δ*hfq* cells. This is consistent with our previous observation that the global regulatory state of the host can modulate phage lytic development, as shown for P1vir infection of an *E. coli* Δ*dksA* mutant (50).

#### Host program in response to P1vir infection

P1vir infection triggers a broad, time-dependent reprogramming in wild-type cells with a progressive increase in transcriptional responsiveness during infection, whereas the Δ*hfq* mutant shows a much narrower, constrained response. In wild-type cells, P1vir infection elicited a strong induction of central carbon and amino acid metabolism (PTS, glycolysis/TCA, PPP), consistent with the metabolic reallocation for other lytic phage-host systems (54–57). Repression of ribosomes, RNA degradation, and nucleotide metabolism can stabilize specific transcripts and rebalance translational capacity in ways that favor phage gene expression, in line with transcriptome-wide studies of phage-induced host remodeling (57,58).

Furthermore, late repression of peptidoglycan biosynthesis indicates a strategic diversion of resources away from cell wall building, which may influence susceptibility to lysis at the end of the phage cycle. Overall, the wild-type host is forced to redirect energy and resources toward phage production and lysis.

In contrast, Δ*hfq* shows an early bias toward oxidative phosphorylation and ABC transporters without broad anabolic activation, suggesting limited mobilization of the host cell for phage production. Late strong induction of chaperones (e.g., *dnaK*/*groL*/*S*, *ibpA*, *hslU*/*V*) and proteostasis suggests that the Δ*hfq* host may be dealing with the accumulation of misfolded proteins or stalled assembly intermediates (due to a defective phage morphogenetic program) consistent with the pleiotropic stress phenotypes and altered proteostasis associated with loss of Hfq (17,59–61).

The ribosome is less remodeled, affecting translation reprogramming in the Δ*hfq* host. The Δ*hfq* host does not engage in full metabolic rewiring; instead, it reacts with a stress and protein-refolding response.

#### Phage–host interaction model in the absence of Hfq

Integration of phage and host transcriptomics shows that P1vir phage can efficiently mobilize metabolism and adjust translation in the wild-type host to support its own development, which aligns with robust phage replication, assembly, and progeny release, yielding a high burst. In contrast, the Δ*hfq* mutant enters P1vir infection with a pre-rewired and stress-prone baseline physiology. Previous studies have shown that *E. coli hfq* mutants exhibit slower growth, reduced motility and biofilm formation, and increased sensitivity to oxidative, osmotic, and temperature stresses (9,17,62). The impaired growth of the *hfq* mutants on multiple carbon sources and their altered stress responses indicate that Hfq broadly modulates cellular carbon and nutrient metabolism (62–64). This likely contributes to the blunted host reprogramming and impaired P1vir development observed in the Δ*hfq* strain. P1vir cannot sufficiently reprogram the pre-rewired metabolism of the Δ*hfq* host to sustain efficient phage production. Consequently, such a narrow host induction program can lead to mis-timed phage regulation and deficits in replication, processing, and morphogenesis, resulting in smaller plaques and lower burst in Δ*hfq* cells.

Hfq is known to play key roles in nucleic acid metabolism. It acts as an RNA chaperone that facilitates sRNA–mRNA pairing and modulates translation and RNA turnover (8,10,64). It also binds DNA, alters its topology, and increases its compaction, behaving as nucleoid-associated protein (14,22). Moreover, Hfq modulates DNA transactions, such as replication and recombination; for example, Hfq deficiency affects ColE1-like plasmid replication in *E. coli* (22). Our data are compatible with both direct Hfq–RNA effects (on specific phage transcripts) and Hfq–DNA interactions, as well as indirect effects via altered host physiology. On the one hand, transcription of specific phage genes may be increased when the phage genome is more relaxed in the absence of Hfq, so that particular phage transcripts are produced prematurely. On the other hand, the loss of Hfq may affect translation or mRNA decay (by either repressing or accelerating degradation), leading to changes in the relative abundance of specific phage transcripts.

Hfq regulates the fate of mRNAs and sRNAs by modulating translation and RNA stability. Hfq can either block or expose ribosome-binding sites (RBS) to repress or activate translation and can protect RNAs from degradation or, conversely, promote their decay via sRNA–mRNA pairing and polyadenylation-coupled turnover, in part through its interactions with RNase E or poly(A) polymerase I (PAP I) (12,65,66). Consistent with possible direct Hfq–RNA interactions, several strongly under-induced P1vir transcripts carry Hfq-favored sequence features (5’ AAN/ARN clusters or 3′ poly-U tracts). Although these motif-based analyses are underpowered by the small number of P1 genes and simple pattern definitions, experimental validation of these candidate P1 transcripts would verify the direct Hfq–RNA interaction hypothesis in these cases. Taken together with our motif scan, these results support a model in which Hfq may directly modulate a subset of P1vir transcripts.

#### Limitations of the study

Our conclusions are based on short-interval RNA-seq (10/20 min) in a P1vir background that is constitutively lytic; thus, we cannot resolve finer timing transitions, later infection stages (e.g., packaging and lysis), or the lysogenic cycle. Adding a denser time course with time-matched mock controls would allow for a more precise mapping of the early → late transition. We could not infer direct Hfq targets from transcript levels alone. RNA-seq also conflates transcription and decay, and mRNA changes may not reflect protein abundance or activity; therefore, proteomic or enzymatic readouts would increase confidence. Targeted follow-ups may also include testing transcript stability and translation, complementation with wild-type or binding-defective Hfq variants, or targeted binding assays for mapping direct Hfq–phage RNA interactions.

## 5. Conclusions

Proper management of bacterial energy and resources and regulation of the phage developmental program over time are key to a successful phage life cycle. It seems evolutionarily justified that control over the cell occurs through its global regulators. Our results identified Hfq as a key hub that enables coordinated host reprogramming and proper timing of the P1vir lytic program in *E. coli*. In its absence, both host and phage responses are impaired, leading to poorly productive interactions.

### Supplementary Materials

**Table S1.** Differential expression of P1vir genes (CSV file); **Table S2.** Lists of overlapping phage genes (in Supplementary Materials file); **Table S3.** Differential expression of host genes (CSV file); **Table S4.** Lists of overlapping host genes (in Supplementary Materials file); **Table S5.** Top ten differentially expressed host genes following P1vir infection (in Supplementary Materials file); **Table S6.** KEGG pathway enrichment results (CSV file); **Table S7.** GO term enrichment results (CSV file); **Table S8.** Hfq-favored motif scan of P1vir transcripts (CSV file); **Table S9.** Statistics for candidate Hfq-sensitive motifs in P1vir transcripts (in the Supplementary Materials file). For CSV files, table captions and column descriptions are provided in the Supplementary Materials file. **Script 1.** P1 motif scan (PY file); **Script 2.** P1 motif stats (PY file).

## Supporting information

Supplementary Materials, Tables, Scripts

## Author Contributions

Conceptualization: GC, AK, ASP; Software: GC, JG, AK; Investigation: GC, JG, AK; Writing—original draft preparation: GC, AK; Writing—review and editing: GC, AK, ASP, GW; Visualization: GC; Supervision: GC; Project administration: GC, GW; Funding acquisition: GC, GW. All authors have read and agreed to the published version of this manuscript.

## Acknowledgments

We thank Magdalena Narajczyk and Sebastian Chamera for their assistance in acquiring electron micrographs. During the preparation of this work, the authors used ChatGPT 5.1 (OpenAI, San Francisco, CA, USA) and Paperpal version 4.4.6 (Cactus Communications Services Pte. Ltd., Singapore) to copyedit English, search the literature, and generate the code. After using these tools and services, the authors reviewed and edited the content as needed and take full responsibility for the content of the published work.

## Institutional Review Board Statement

Not applicable.

## Informed Consent Statement

Not applicable.

## Data Availability Statement

RNA sequencing data were deposited in the NCBIʹs Gene Expression Omnibus (GEO) and are accessible through the GEO Series accession number GSE262812.

