## Supplementary Materials, Tables, Scripts for "Virus-Host Interaction Gets *Lousy*: P1vir Phage Development Upon Impaired RNA Global Regulation in *Escherichia coli* Δ*hfq* Mutant Cells": Script1_p1_motif_scan.py

```
#!/usr/bin/env python
"""
Scan P1 phage genes for Hfq-favoured motifs (AAN / ARN in 5' windows,
poly-U in 3' windows), and (optionally) merge with DE data.

Windows (nt) relative to CDS, oriented to mRNA sense:
    W5: -30..+60 around the start codon
    W3: -40..+20 around the stop codon

NEW:
  - Also report AAN_max_run and ARN_max_run (max cluster length in triplets).
  - Scan ALL CDS genes from the GBFF; attach ddlog2FC / IRD only if present
    in the DE CSV (otherwise leave as NaN).
"""

import sys
import re
import pandas as pd
from Bio import SeqIO
from Bio.Seq import Seq

# Windows (nt) relative to CDS, oriented to mRNA sense
W5_UP, W5_DN = 30, 60   # 5′ window: -30..+60 around start codon
W3_UP, W3_DN = 40, 20   # 3′ window: -40..+20 around stop codon


def load_genes_from_gbff(gbff_path):
    """
    Parse GenBank (GBFF) and return:
    - genome_seq: full sequence (assumes single main record)
    - gene_info: dict gene_name -> dict(start, end, strand)
      coordinates are 0-based, end is exclusive, as in Biopython.
    """
    records = list(SeqIO.parse(gbff_path, "genbank"))
    if len(records) == 0:
        raise ValueError(f"No records found in {gbff_path}")
    if len(records) > 1:
        print(f"[WARN] Multiple records found in {gbff_path}; using the first one: {records[0].id}", file=sys.stderr)

    rec = records[0]
    genome_seq = rec.seq
    gene_info = {}

    for feat in rec.features:
        if feat.type != "CDS":
            continue

        qualifiers = feat.qualifiers
        gene_name = None

        # Prefer "gene", fall back to "locus_tag"
        if "gene" in qualifiers:
            gene_name = qualifiers["gene"][0]
        elif "locus_tag" in qualifiers:
            gene_name = qualifiers["locus_tag"][0]

        if not gene_name:
            continue

        start = int(feat.location.start)
        end = int(feat.location.end)
        strand = int(feat.location.strand)  # +1 or -1

        gene_info[gene_name] = {
            "start": start,
            "end": end,
            "strand": strand
        }

    return genome_seq, gene_info


def get_windows(genome_seq, start, end, strand):
    """
    Extract 5' (W5) and 3' (W3) windows in transcript orientation.

    Definitions (relative to CDS boundaries, approximating start/stop codons):
      - W5: -30..+60 nt around the start codon
      - W3: -40..+20 nt around the stop codon

    For + strand:
        start codon ~ CDS.start
        stop codon  ~ CDS.end
        W5 genomic   = [start - W5_UP, start + W5_DN)
        W3 genomic   = [end   - W3_UP, end   + W3_DN)

    For - strand:
        transcript runs from genomic end -> start.
        In genomic coordinates:
        - start codon (5' in mRNA) ~ CDS.end
        - stop codon  (3' in mRNA) ~ CDS.start
        So:
        W5 genomic   = [end   - W5_DN, end   + W5_UP)
        W3 genomic   = [start - W3_DN, start + W3_UP)
        Then both windows are reverse-complemented to get mRNA sense.
    """
    seqlen = len(genome_seq)

    if strand == 1:
        # 5' window: -30..+60 around start
        w5_start = max(0, start - W5_UP)
        w5_end = min(seqlen, start + W5_DN)
        w5 = genome_seq[w5_start:w5_end]

        # 3' window: -40..+20 around stop
        w3_start = max(0, end - W3_UP)
        w3_end = min(seqlen, end + W3_DN)
        w3 = genome_seq[w3_start:w3_end]

        # Already in transcript orientation
        w5_seq = w5.upper()
        w3_seq = w3.upper()

    elif strand == -1:
        # 5' window in mRNA: -30..+60 around start codon at genomic 'end'
        w5_start = max(0, end - W5_DN)
        w5_end = min(seqlen, end + W5_UP)
        w5 = genome_seq[w5_start:w5_end]

        # 3' window in mRNA: -40..+20 around stop codon at genomic 'start'
        w3_start = max(0, start - W3_DN)
        w3_end = min(seqlen, start + W3_UP)
        w3 = genome_seq[w3_start:w3_end]

        # Reverse complement to get transcript orientation
        w5_seq = w5.reverse_complement().upper()
        w3_seq = w3.reverse_complement().upper()
    else:
        raise ValueError(f"Unexpected strand value: {strand}")

    return str(w5_seq), str(w3_seq)


def count_AAN_ARN_runs(seq5):
    """
    Count AAN and ARN clusters in a 5' window and get max run lengths (in triplets).

    We define:
      - AAN triplet: AA[ACGT]
      - ARN triplet: A[AG][ACGT]  (R = A or G)
    A cluster is >= 3 consecutive triplets of that type.
    We return:
      AAN_repeats, AAN_max_run, ARN_repeats, ARN_max_run
    """
    seq = seq5.upper()

    # Patterns: 3 or more consecutive triplets
    AAN_pat = re.compile(r'(?:AA[ACGT]){3,}')
    ARN_pat = re.compile(r'(?:A[AG][ACGT]){3,}')

    def _count_and_max(pattern):
        matches = list(pattern.finditer(seq))
        if not matches:
            return 0, 0
        count = len(matches)
        max_run = 0
        for m in matches:
            nt_len = m.end() - m.start()
            triplets = nt_len // 3
            if triplets > max_run:
                max_run = triplets
        return count, max_run

    aan_count, aan_max = _count_and_max(AAN_pat)
    arn_count, arn_max = _count_and_max(ARN_pat)

    return aan_count, aan_max, arn_count, arn_max


def count_polyU_features(seq3):
    """
    In the 3' window, look for poly-U–like features and UA repeats.
    We work on an RNA version (T->U).
    Returns:
      UA_repeats      = number of UA di-nucleotide runs (>=2 consecutive "UA")
      UA_max_repeats  = max length (in UA units) of such runs
      polyU_count     = number of U-runs (>=3 U's)
      polyU_max       = max length (in U's) of such runs
    """
    rna = seq3.upper().replace("T", "U")

    # UA runs: (UA){2,} etc.
    UA_pat = re.compile(r'(?:UA){2,}')
    ua_matches = list(UA_pat.finditer(rna))
    ua_repeats = len(ua_matches)
    ua_max = 0
    for m in ua_matches:
        run_len = (m.end() - m.start()) // 2  # length in UA repeats
        if run_len > ua_max:
            ua_max = run_len

    # Poly-U runs: U{3,}
    polyU_pat = re.compile(r'U{3,}')
    poly_matches = list(polyU_pat.finditer(rna))
    polyU_count = len(poly_matches)
    polyU_max = 0
    for m in poly_matches:
        run_len = m.end() - m.start()  # length in U's
        if run_len > polyU_max:
            polyU_max = run_len

    return ua_repeats, ua_max, polyU_count, polyU_max


def main(gbff_path, de_csv_path, out_csv_path):
    # Load genome and gene coordinates
    genome_seq, gene_info = load_genes_from_gbff(gbff_path)

    # DE table is optional: if path == "-" treat as no DE file
    if de_csv_path != "-":
        de = pd.read_csv(de_csv_path)
        if "Gene" not in de.columns:
            raise ValueError("Input DE table must contain a 'Gene' column.")
        de = de.set_index("Gene")
    else:
        de = None

    rows = []

    # IMPORTANT: iterate over ALL genes from GBFF
    for gene, info in gene_info.items():
        start = info["start"]
        end = info["end"]
        strand = info["strand"]

        w5_seq, w3_seq = get_windows(genome_seq, start, end, strand)
        w5_len = len(w5_seq)
        w3_len = len(w3_seq)

        # Motif features
        AAN_repeats, AAN_max_run, ARN_repeats, ARN_max_run = count_AAN_ARN_runs(w5_seq)
        UA_repeats, UA_max_repeats, polyU_count, polyU_max = count_polyU_features(w3_seq)

        # Default DE metrics = NaN
        dd10 = dd20 = ird10 = ird20 = None

        # If DE table is provided and gene exists there, pull the values
        if de is not None and gene in de.index:
            if "ddlog2FC_10" in de.columns:
                dd10 = de.at[gene, "ddlog2FC_10"]
            if "ddlog2FC_20" in de.columns:
                dd20 = de.at[gene, "ddlog2FC_20"]
            if "IRD_10" in de.columns:
                ird10 = de.at[gene, "IRD_10"]
            if "IRD_20" in de.columns:
                ird20 = de.at[gene, "IRD_20"]

        # Rank score: favour strong under-induction and strong motifs
        rank_score = None
        if dd20 is not None and pd.notnull(dd20):
            rank_score = (-dd20) + 0.3 * AAN_repeats + 0.3 * ARN_repeats + 0.2 * polyU_max

        rows.append(
            {
                "Gene": gene,
                "w5_len": w5_len,
                "w3_len": w3_len,
                "AAN_repeats": AAN_repeats,
                "AAN_max_run": AAN_max_run,
                "ARN_repeats": ARN_repeats,
                "ARN_max_run": ARN_max_run,
                "UA_repeats": UA_repeats,
                "UA_max_repeats": UA_max_repeats,
                "polyU_count": polyU_count,
                "polyU_max": polyU_max,
                "ddlog2FC_10": dd10,
                "ddlog2FC_20": dd20,
                "IRD_10": ird10,
                "IRD_20": ird20,
                "rank_score": rank_score,
            }
        )

    out_df = pd.DataFrame(rows)
    out_df.to_csv(out_csv_path, index=False)
    print(f"[INFO] Wrote motif scan results for {len(out_df)} genes to {out_csv_path}")


if __name__ == "__main__":
    if len(sys.argv) != 3 and len(sys.argv) != 4:
        print(
            "Usage:\n"
            "  python p1_motif_scan.py P1.gbff phage_DE_table.csv [P1_motif_scan_results.csv]\n"
            "  (use '-' instead of phage_DE_table.csv if you have no DE data)",
            file=sys.stderr,
        )
        sys.exit(1)

    gbff_file = sys.argv[1]
    de_file = sys.argv[2]
    out_file = sys.argv[3] if len(sys.argv) == 4 else "P1_motif_scan_results.csv"
    main(gbff_file, de_file, out_file)
```
