## Supplementary Materials, Tables, Scripts for "Virus-Host Interaction Gets *Lousy*: P1vir Phage Development Upon Impaired RNA Global Regulation in *Escherichia coli* Δ*hfq* Mutant Cells": Script2_p1_motif_stats.py

```
#!/usr/bin/env python
"""
Test enrichment of stronger under-induction among P1 genes
with Hfq-favoured motif features.

Input:  P1_motif_scan_results.csv  (from p1_motif_scan.py)
        must contain columns:
        Gene, AAN_max_run, ARN_max_run, polyU_max, UA_max_repeats,
        ddlog2FC_10, ddlog2FC_20, IRD_10, IRD_20

Output: P1_motif_enrichment_summary.csv
"""

import sys
import numpy as np
import pandas as pd
from scipy.stats import fisher_exact, mannwhitneyu


# -----------------------------
# Parameters
# -----------------------------

# Motif thresholds
AAN_RUN_MIN = 3      # at least 3 AAN triplets in a cluster
ARN_RUN_MIN = 3      # at least 3 ARN triplets in a cluster
UA_RUN_MIN  = 3      # at least 3 UA in a cluster (adjust as needed)
POLYU_MIN   = 5      # at least 5 U in a run at 3'

# “Strong under-induction” by IRD (hfq vs WT)
IRD_LOWER = 0.8      # IRD <= 0.8 considered under-induced


def main(motif_csv, out_csv="P1_motif_enrichment_summary.csv"):
    df = pd.read_csv(motif_csv)

    # --- Basic cleaning ---
    # ensure numeric
    for col in ["AAN_max_run", "ARN_max_run", "polyU_max", "UA_max_repeats",
                "ddlog2FC_10", "ddlog2FC_20", "IRD_10", "IRD_20"]:
        if col in df.columns:
            df[col] = pd.to_numeric(df[col], errors="coerce")

    # -----------------------------
    # Define motif presence flags
    # -----------------------------
    df["motif_AAN"]   = df["AAN_max_run"] >= AAN_RUN_MIN
    df["motif_ARN"]   = df["ARN_max_run"] >= ARN_RUN_MIN
    df["motif_polyU"] = df["polyU_max"]   >= POLYU_MIN
    df["motif_UA"]    = df["UA_max_repeats"]      >= UA_RUN_MIN
    df["motif_any"]   = df[["motif_AAN", "motif_ARN", "motif_polyU", "motif_UA"]].any(axis=1)

    # ----------------------------------------
    # Define "strong under-induction" (any time)
    # ----------------------------------------
    df["strong_down_10"] = df["IRD_10"] <= IRD_LOWER
    df["strong_down_20"] = df["IRD_20"] <= IRD_LOWER
    df["strong_down_any"] = df[["strong_down_10", "strong_down_20"]].any(axis=1)

    # -----------------------------
    # Helper: MWU for a given ddlog2FC column
    # -----------------------------
    def mwu_for_col(present, absent, col_name):
        """Return (stat, p, median_present, median_absent) for given ddlog2FC column."""
        if col_name not in df.columns:
            return np.nan, np.nan, np.nan, np.nan

        dd_present = present[col_name].dropna()
        dd_absent  = absent[col_name].dropna()

        if len(dd_present) == 0 or len(dd_absent) == 0:
            return np.nan, np.nan, np.nan, np.nan

        try:
            stat, pval = mannwhitneyu(dd_present, dd_absent, alternative="two-sided")
        except Exception:
            stat, pval = np.nan, np.nan

        median_present = dd_present.median()
        median_absent  = dd_absent.median()

        return stat, pval, median_present, median_absent

    # -----------------------------
    # Helper: Fisher + MWU per motif
    # -----------------------------
    def analyze_motif(flag_col):
        sub = df.dropna(subset=["strong_down_any"])  # ensure boolean is valid

        # 2x2 table for Fisher: motif present/absent vs strong_down_any
        present = sub[sub[flag_col]]
        absent  = sub[~sub[flag_col]]

        a = present["strong_down_any"].sum()            # motif+, strong_down+
        b = present["strong_down_any"].size - a         # motif+, strong_down-
        c = absent["strong_down_any"].sum()             # motif-, strong_down+
        d = absent["strong_down_any"].size - c          # motif-, strong_down-

        table = np.array([[a, b],
                          [c, d]], dtype=float)

        # Fisher’s exact (odds ratio >1 means enrichment of strong_down in motif+)
        try:
            OR, p_fisher = fisher_exact(table)
        except Exception:
            OR, p_fisher = np.nan, np.nan

        # Mann–Whitney for both ddlog2FC_20 and ddlog2FC_10
        mwu20_stat, mwu20_p, med20_plus, med20_minus = mwu_for_col(
            present, absent, "ddlog2FC_20"
        )
        mwu10_stat, mwu10_p, med10_plus, med10_minus = mwu_for_col(
            present, absent, "ddlog2FC_10"
        )

        return {
            "motif_flag": flag_col,
            "n_motif_plus": int(sub[flag_col].sum()),
            "n_motif_minus": int((~sub[flag_col]).sum()),
            "a_motif+_strong+": int(a),
            "b_motif+_strong-": int(b),
            "c_motif-_strong+": int(c),
            "d_motif-_strong-": int(d),
            "fisher_OR": OR,
            "fisher_p": p_fisher,

            # ddlog2FC_10 MWU + medians
            "MWU_stat_ddlog2FC_10": mwu10_stat,
            "MWU_p_ddlog2FC_10": mwu10_p,
            "ddlog2FC_10_median_motif+": med10_plus,
            "ddlog2FC_10_median_motif-": med10_minus,
            
            # ddlog2FC_20 MWU + medians
            "MWU_stat_ddlog2FC_20": mwu20_stat,
            "MWU_p_ddlog2FC_20": mwu20_p,
            "ddlog2FC_20_median_motif+": med20_plus,
            "ddlog2FC_20_median_motif-": med20_minus,
        }

    # Run for each motif class (including UA)
    motif_cols = ["motif_AAN", "motif_ARN", "motif_polyU", "motif_UA", "motif_any"]
    results = [analyze_motif(col) for col in motif_cols]

    out = pd.DataFrame(results)
    out.to_csv(out_csv, index=False)
    print(f"[INFO] Wrote motif enrichment summary to {out_csv}")


if __name__ == "__main__":
    if len(sys.argv) < 2:
        print("Usage: python p1_motif_stats.py P1_motif_scan_results.csv [output.csv]", file=sys.stderr)
        sys.exit(1)

    motif_csv = sys.argv[1]
    out_csv = sys.argv[2] if len(sys.argv) >= 3 else "P1_motif_enrichment_summary.csv"
    main(motif_csv, out_csv)
```
