## Supplementary Materials, Tables, Scripts for "Virus-Host Interaction Gets *Lousy*: P1vir Phage Development Upon Impaired RNA Global Regulation in *Escherichia coli* Δ*hfq* Mutant Cells": Supplementary Materials.pdf

### Table S1. Differential expression of P1vir genes (CSV file).

Phage transcriptome from infections of wild-type and Δ*hfq* hosts was assessed using RNA-seq. We assessed gene expression at 10 min (early stage) and 20 min (late stage) post-infection. For each gene, the log2 fold change (log2FC) in expression was calculated relative to the no-phage sample (indicated in this study as infection time 0 min). WT, wild type; hfq, Δ*hfq* mutant.

Column descriptions:

- No – consecutive row number.
- Locus tag – annotated locus identifier of the phage gene.
- GeneID – unique identifier assigned by NCBI database.
- Gene symbol – gene name as given in the annotation (systematic name) and used throughout the text/figures.
- Strand – DNA strand on which the gene is encoded (+ or −).
- Gene description (NCBI) – gene description obtained from the NCBI Gene database.
- Function description – brief functional description of the gene product according to Łobocka et al. (2004), with updates (Piya et al. 2018, Bednarek et al. 2022, Zhao et al. 2025).
- Module – functional category assigned to the gene in this study.
- log2FC_WT_0 – $\log_2$ fold change in phage gene expression at 0 min vs. no phage in the wild-type host.
- padj_WT_0 – adjasted p-value calculated for log2FC_WT_0.
- log2FC_hfq_0 – $\log_2$ fold change in phage gene expression at 0 min vs. no phage in the Δ*hfq* host.
- padj_hfq_0 – adjasted p-value calculated for log2FC_hfq_0.
- log2FC_WT_10 – $\log_2$ fold change in phage gene expression at 10 min vs. 0 min (= no phage) in the wild-type host.
- padj_WT_10 – adjasted p-value calculated for log2FC_WT_10.
- log2FC_hfq_10 – $\log_2$ fold change in phage gene expression at 10 min vs. 0 min (= no phage) in the Δ*hfq* host.
- padj_hfq_10 – adjasted p-value calculated for log2FC_hfq_10.

- ddlog2FC_10 – between-strain contrast at 10 min: ($log_2FC_{\Delta hfq} - log_2FC_{WT}$).
- IRD_10 – induction ratio difference (Δ*hfq* vs. WT) at 10 min, calculated as 2^(ddlog2FC_10).
- sig10 – significance flag for differential expression at 10 min; WT or hfq indicates that the gene is significantly differentially expressed vs. 0 min (FDR < 0.05) in the respective host at this time point.
- log2FC_WT_20 – $log_2$ fold change in phage gene expression at 20 min vs. 0 min (= no phage) in the wild-type host.
- padj_WT_20 – adjasted p-value calculated for log2FC_WT_20.
- log2FC_hfq_20 – $log_2$ fold change in phage gene expression at 10 min vs. 0 min (= no phage) in the Δ*hfq* host.
- padj_hfq_20 – adjasted p-value calculated for log2FC_hfq_20.
- ddlog2FC_20 – between-strain contrast at 20 min: ($log_2FC_{\Delta hfq} - log_2FC_{WT}$).
- IRD_20 – induction ratio difference (Δ*hfq* vs. WT) at 20 min, calculated as 2^(ddlog2FC_20).
- sig20 – significance flag for differential expression at 20 min; WT or hfq indicates that the gene is significantly differentially expressed vs. 0 min (FDR < 0.05) in the respective host at this time point.

**Table S2. Lists of overlapping phage genes.** Gene lists corresponding to Venn diagram showing the overlap (intersections) upregulated phage genes between hosts and time points. Abbreviations: 10, 10 min post-infection; 20, 20 min post-infection; wt, wild type; hfq, Δ*hfq* mutant.

| Intersections | Genes |
|---|---|
| WT_10 ∩ WT_20 ∩ hfq_10 | cra |
| WT_10 ∩ WT_20 ∩ hfq_10 ∩ hfq_20 | cre, c8, ref, ssb, isaA, icd, ant1, kilA, repL, parB, parA, uhr, hrdC, dmt, plp, tciA, tciB, tciC, trnI, ban, pmgL, pmgM, pmgN, pmgO, pmgP, ppp, pmgQ, pmgR, pmgS, pap, pmgT, pmgU, pmgV, pdcB, c1 |
| WT_10 ∩ WT_20 ∩ hfq_20 | c4 |
| WT_10 ∩ WT_20 | doc |
| WT_20 ∩ hfq_20 | rlfA, trnA, upfM |
| WT_20 | rlfB |
| WT_20 ∩ hfq_10 ∩ hfq_20 | upl, lpa, coi |
| hfq_20 | dbn |

**Table S3. Differential expression of host genes (CSV file).**

Host transcriptomes from P1vir infections of the wild-type and Δ*hfq* strains were analyzed using RNA-seq. Gene expression was assessed at 10 min (early stage) and 20 min (late stage) post-infection. For each gene, the log2 fold change (log2FC) in expression was calculated relative to the no-phage sample (indicated in this study as infection time 0 min). WT, wild type; hfq, Δ*hfq* mutant.

Column descriptions:

- STRAIN – indicates host strain, time point, and regulation in expression calculated relative vs. no phage transcriptome at a given time point (e.g., HOST_DE_P1_10_vs_nophage__hfq_downregulated = host differential expression upon P1vir infection at 10 min vs. 0 min (= no phage) control in Δ*hfq* strain, downregulated genes).
- Locus tag – annotated locus identifier of the *E. coli* gene.
- GeneID – unique identifier assigned by NCBI database.
- Fold change – $\log_2$ fold change in host gene expression vs. 0 min (= no phage).
- Adjusted p-value – significance level taking into account the false discovery rate (FDR).
- Gene symbol – gene name as given in the annotation (systematic name) and used throughout the text/figures.
- Biotype – properties of a gene; possible values are tRNA, rRNA, snRNA, scRNA, snoRNA, miscRNA, ncRNA, protein coding, pseudo, other, and unknown.
- Description – gene product function.

**Table S4. Lists of overlapping host genes.** Gene lists corresponding to Venn diagrams showing the overlap (intersections) of downregulated and upregulated genes between hosts and time points. Abbreviations: 10, 10 min post-infection; 20, 20 min post-infection; wt, wild type; hfq, Δ*hfq* mutant; down, downregulated; up, upregulated.

| Intersections | Genes |
|---|---|
| 10_wt_down ∩ 20_wt_down ∩ 10_hfq_down ∩ 20_hfq_down | cspG, cspA, glpD, rho, nusA, |
| 10_wt_down ∩ 20_wt_down | entC, ylaB, sthA, cycA, mtlA, yhjV, cirA, fecI, entE, kgtP, fhuF, ybiP, yidD, ampG, fimA, rsmB, obgE, rnpA, murC, mrdA, gtrB, pdhR, yhbE, exbB, rrlB, rrlH, mnmG |
| 10_wt_down ∩ 20_hfq_down | yfiM |
| 10_wt_down ∩ 20_wt_down ∩ 10_hfq_down | yncD, dinG, rimP |
| 10_wt_down | ybiR, proP, efeB, rseB, efeO, yoaE, wzxB, rrlD, rrlE, rrlG |
| 10_wt_down ∩ 20_wt_down ∩ 20_hfq_down | nlpI, deaD, rhlE, suhB |
| 20_wt_down | ydjN, fecR, opgC, entA, rsxD, fhuA, intB, entB, prmC, nth, fepB, metY, ydgI, ydiU, yfiB, fhuB, yrbN, exbD, queD, fiu, waaS, rlmA, rluE, wecD, mreD, rlmH, yegD, rsmG, carA, wzxE, queA, rnd, yehS, yojI, epmA, lolE, yabP, fadR, hha, dusA, rluA, prfA, infA, yraQ, yhiN, der, rlmG, fhuD, truB, panF, rapA, dusB, mdfA, mnmE, ygiQ, lpxH, pnp, msbA, rbfA, tonB, yncE, pyrD, srmB, lpxB, rluB, dnaE, waaL, ndk, gpt, codA, pitA, yccA, udk, rnc, lysW, ycbZ, era, rrsB, trmA, yidC, typA, oppD, uvrC, holA, secF, bax, wzzE, mreC, oppC, recF, uup, rimO, prc, eptB, rlmN, tgt, cdsA, dnaX, ybiT, rsxC, gmk, mrcA, ldtB, purR, potA, yejM, potB, oppF, ddlB, rluC, rlmL, mltD, rrlA, yifK, aceE, mnmA, ispU, lpd, hpt, cmk, rpoB, dnaA, wecC, lysT, ycaO, flu, rpsH, plaP, rrsG, ispE, accB, pheT, oppA, rplX, plsX, pyrH, fadL, fecA, spoT, fis, rhlB, prfC, secY, trmD, rplN, rpsJ, accC, infC, rpmE, rplI, rplE, rplS, rpsF, pyrG, rplR, lepA, rplA, lpxC, rplD, gltX, rplM, priB, rimM, rpsA, rplC, rpoA, fabH, rpmB, proQ, rpsL, rpsM, rpsD, rpoC, rpmH, rplO, rpsG, rplK, rpmJ, rplL, tsf, rpsP, rpsI, fusA, rpmI, rplJ, tufA, rpsB, rpsS |
| 20_wt_down ∩ 20_hfq_down | infB, gltT, rplF, aceF, rpsO, rpsE, rpsK, cyoA, rplB |
| 20_wt_down ∩ 10_hfq_down ∩ 20_hfq_down | ydfJ, rnr |
| 10_hfq_down | yejG, insJ, bluF, insK, yzgL |

| | |
|---|---|
| 20_hfq_down | lpxP, glgP, nsrR |
| 10_wt_up | cysJ, cysK, glnP, ybjN, fruA, ydjN, dhaM, ybgE, thrT, speB, speA, sdaC, upp, gyrA, hns, atpG, ptsG, tig, atpD, tktA, rplJ |
| 10_wt_up ∩ 20_wt_up ∩ 10_hfq_up ∩ 20_hfq_up | glnH |
| 10_wt_up ∩ 20_wt_up | gatY, glnQ, manZ, yjiY, pflA, fruB, deoC, asnA, fruK, cydA, gatZ, uspA, thrA, manX, cydB, lpp, cspC, malT, pmbA, ppc, ompF, crp, trpS, pgk, rnpB, kbl, dapA, recA, arcA, sucA, gapA, clpX, asnS, ileS |
| 10_wt_up ∩ 20_wt_up ∩ 20_hfq_up | glnA, adhE, pntA, tufB |
| 10_wt_up ∩ 10_hfq_up | metN, metI, metK, iscS |
| 10_wt_up ∩ 20_wt_up ∩ 10_hfq_up | glyA |
| 20_wt_up | pstC, alaC, gatA, pstS, yagU, ibpB, ycjX, gatC, pstB, yghJ, phoU, ybgK, yhiI, ldhA, pntB, udp, pgl, thiC, manY, hinT, umuC, mlc, glgC, glgB, sucC, slyB, yeaD, yhaM, uxuB, uxuA, glgX, moaE, glgA, lon, yfbU, yqjI, ydfG, nupG, dnaJ, ghrB, ybeZ, cdd, gltB, frdA, mdh, thrC, ybgI, ydhQ, fbp, lrp, sodB, ushA, sucB, serC, gcvP, gltD, hslO, tas, dadA, nagE, sucD, ribC, lysU, metH, asd, rpoS, aroA, panD, gltA, galM, mscS, tyrB, yjhU, dcuA, ompR, pykA, hupA, yhgF, glnE, yobF, oxyR, clpP, proB, yqhD, hsdR, nadE, minD, nfsB, pfkA, frmA, pfo, tpiA, pepA, dksA, zwf, uvrA, lexA, nuoF, galU, gnd, yajQ, pepN, hupB, serS, dapD, deoD, pepD, pgi |
| 20_wt_up ∩ 20_hfq_up | clpB, ibpA, groS, dnaK, groL, recN, ybbN, htpG, hslV, prlC, hslU, grpE |
| 20_wt_up ∩ 10_hfq_up ∩ 20_hfq_up | glmS |
| 20_wt_up ∩ 10_hfq_up | atpE |
| 10_hfq_up | soxS, fxsA, iscR, metQ, iscA, atpF, dacA, plaP, hscA, atpA |
| 10_hfq_up ∩ 20_hfq_up | iscU |
| 20_hfq_up | hslR, dsbA |

**Table S5. Top ten differentially expressed host genes following P1*vir* infection.** The table lists the ten most downregulated (▼) and ten most upregulated (▲) host genes at each time point (10 and 20 min post-infection) for both Δ*hfq* and wild-type strains. Fold change is reported relative to 0 min (= no phage) and expressed as log2FC. Gene symbols and brief descriptions are provided. Abbreviations: Reg, regulation; FC, fold change; WT, wild-type.

| HOST | Reg | Time | TOP | FC | Gene | Description |
|---|---|---|---|---|---|---|
| Δ*hfq* | ▼ | 10 | **1** | 4.1 | ***cspA*** | RNA chaperone and antiterminator, cold-inducible |
| Δ*hfq* | ▼ | 10 | **2** | 3.6 | ***cspG*** | cold shock protein homolog, cold-inducible |
| Δ*hfq* | ▼ | 10 | **3** | 2.8 | ***yejG*** | uncharacterized protein |
| Δ*hfq* | ▼ | 10 | **4** | 2.6 | ***insJ*** | IS150 transposase A |
| Δ*hfq* | ▼ | 10 | **5** | 2.6 | ***bluF*** | anti-repressor for YcgE, blue light-responsive; FAD-binding; has c-di-GMP phosphodiesterase-like EAL domain, but does not degrade c-di-GMP |
| Δ*hfq* | ▼ | 10 | **6** | 2.4 | ***insK*** | IS150 transposase B |
| Δ*hfq* | ▼ | 10 | **7** | 2.3 | ***yzgL*** | pseudogene |
| Δ*hfq* | ▼ | 10 | **8** | 2.1 | ***yncD*** | putative iron outer membrane transporter |
| Δ*hfq* | ▼ | 10 | **9** | 1.8 | ***dinG*** | ATP-dependent DNA helicase |
| Δ*hfq* | ▼ | 10 | **10** | 1.8 | ***glpD*** | sn-glycerol-3-phosphate dehydrogenase, aerobic, FAD/NAD(P)-binding |
| Δ*hfq* | ▼ | 20 | **1** | 7.4 | ***cspG*** | cold shock protein homolog, cold-inducible |
| Δ*hfq* | ▼ | 20 | **2** | 5.0 | ***cspA*** | RNA chaperone and antiterminator, cold-inducible |
| Δ*hfq* | ▼ | 20 | **3** | 2.5 | ***lpxP*** | palmitoleoyl-acyl carrier protein (ACP)-dependent acyltransferase |
| Δ*hfq* | ▼ | 20 | **4** | 2.3 | ***yfiM*** | putative lipoprotein |
| Δ*hfq* | ▼ | 20 | **5** | 2.1 | ***suhB*** | inositol monophosphatase |
| Δ*hfq* | ▼ | 20 | **6** | 2.0 | ***glgP*** | glycogen phosphorylase |
| Δ*hfq* | ▼ | 20 | **7** | 2.0 | ***nsrR*** | nitric oxide-sensitive repressor for NO regulon |
| Δ*hfq* | ▼ | 20 | **8** | 1.9 | ***glpD*** | sn-glycerol-3-phosphate dehydrogenase, aerobic, FAD/NAD(P)-binding |
| Δ*hfq* | ▼ | 20 | **9** | 1.8 | ***ydfJ*** | pseudogene |
| Δ*hfq* | ▼ | 20 | **10** | 1.8 | ***rnr*** | exoribonuclease R, RNase R |
| Δ*hfq* | ▲ | 10 | **1** | 3.1 | ***soxS*** | superoxide response regulon transcriptional activator; autoregulator |
| Δ*hfq* | ▲ | 10 | **2** | 2.5 | ***fxsA*** | suppressor of F exclusion of phage T7 |
| Δ*hfq* | ▲ | 10 | **3** | 2.3 | ***glnH*** | glutamine transporter subunit |
| Δ*hfq* | ▲ | 10 | **4** | 1.9 | ***metI*** | DL-methionine transporter subunit |
| Δ*hfq* | ▲ | 10 | **5** | 1.8 | ***metK*** | S-adenosylmethionine synthetase |
| Δ*hfq* | ▲ | 10 | **6** | 1.8 | ***metN*** | DL-methionine transporter subunit |

| | | | | | | |
|---|---|---|---|---|---|---|
| Δ*hfq* | ▲ | 10 | **7** | 1.7 | ***iscR*** | isc operon transcriptional repressor; suf operon transcriptional activator; oxidative stress- and iron starvation-inducible; autorepressor |
| Δ*hfq* | ▲ | 10 | **8** | 1.7 | ***iscS*** | cysteine desulfurase (tRNA sulfurtransferase), PLP-dependent |
| Δ*hfq* | ▲ | 10 | **9** | 1.6 | ***glmS*** | L-glutamine:D-fructose-6-phosphate aminotransferase |
| Δ*hfq* | ▲ | 10 | **10** | 1.6 | ***metQ*** | DL-methionine transporter subunit |
| Δ*hfq* | ▲ | 20 | **1** | 3.3 | ***ibpA*** | heat shock chaperone |
| Δ*hfq* | ▲ | 20 | **2** | 3.0 | ***groS*** | Cpn10 chaperonin GroES, small subunit of GroESL |
| Δ*hfq* | ▲ | 20 | **3** | 2.9 | ***groL*** | Cpn60 chaperonin GroEL, large subunit of GroESL |
| Δ*hfq* | ▲ | 20 | **4** | 2.8 | ***dnaK*** | chaperone Hsp70, with co-chaperone DnaJ |
| Δ*hfq* | ▲ | 20 | **5** | 2.5 | ***glnH*** | glutamine transporter subunit |
| Δ*hfq* | ▲ | 20 | **6** | 2.5 | ***hslU*** | molecular chaperone and ATPase component of HslUV protease |
| Δ*hfq* | ▲ | 20 | **7** | 2.3 | ***prlC*** | oligopeptidase A |
| Δ*hfq* | ▲ | 20 | **8** | 2.3 | ***clpB*** | protein disaggregation chaperone |
| Δ*hfq* | ▲ | 20 | **9** | 2.3 | ***grpE*** | heat shock protein |
| Δ*hfq* | ▲ | 20 | **10** | 2.2 | ***hslV*** | peptidase component of the HslUV protease |
| WT | ▼ | 10 | **1** | 7.8 | ***cspG*** | cold shock protein homolog, cold-inducible |
| WT | ▼ | 10 | **2** | 6.9 | ***cspA*** | RNA chaperone and antiterminator, cold-inducible |
| WT | ▼ | 10 | **3** | 2.7 | ***entC*** | isochorismate synthase 1 |
| WT | ▼ | 10 | **4** | 2.7 | ***ylaB*** | putative membrane-anchored cyclic-di-GMP phosphodiesterase |
| WT | ▼ | 10 | **5** | 2.6 | ***glpD*** | sn-glycerol-3-phosphate dehydrogenase, aerobic, FAD/NAD(P)-binding |
| WT | ▼ | 10 | **6** | 2.3 | ***yfiM*** | putative lipoprotein |
| WT | ▼ | 10 | **7** | 2.3 | ***sthA*** | pyridine nucleotide transhydrogenase, soluble |
| WT | ▼ | 10 | **8** | 2.3 | ***yncD*** | putative iron outer membrane transporter |
| WT | ▼ | 10 | **9** | 2.2 | ***cycA*** | D-alanine/D-serine/glycine transporter |
| WT | ▼ | 10 | **10** | 2.1 | ***mtlA*** | fused mannitol-specific PTS enzymes: IIA components/IIB components/IIC components |
| WT | ▼ | 20 | **1** | 8.9 | ***cspG*** | cold shock protein homolog, cold-inducible |
| WT | ▼ | 20 | **2** | 8.0 | ***cspA*** | RNA chaperone and antiterminator, cold-inducible |
| WT | ▼ | 20 | **3** | 4.7 | ***ydjN*** | putative transporter |
| WT | ▼ | 20 | **4** | 3.6 | ***glpD*** | sn-glycerol-3-phosphate dehydrogenase, aerobic, FAD/NAD(P)-binding |
| WT | ▼ | 20 | **5** | 2.9 | ***deaD*** | ATP-dependent RNA helicase |
| WT | ▼ | 20 | **6** | 2.7 | ***fecI*** | RNA polymerase, sigma 19 factor |
| WT | ▼ | 20 | **7** | 2.5 | ***fecR*** | FecI pro-sigma factor; transmembrane signal transducer for ferric citrate transport; periplasmic FecA:ferric citrate sensor and cytoplasmic FecI ECF sigma factor activator |
| WT | ▼ | 20 | **8** | 2.5 | ***suhB*** | inositol monophosphatase |

| | | | | | | |
|---|---|---|---|---|---|---|
| WT | ▼ | 20 | **9** | 2.5 | ***rhlE*** | ATP-dependent RNA helicase |
| WT | ▼ | 20 | **10** | 2.4 | ***cirA*** | colicin IA outer membrane receptor and translocator; ferric iron-catecholate transporter |
| WT | ▲ | 10 | **1** | 4.1 | ***cysJ*** | sulfite reductase, alpha subunit, flavoprotein |
| WT | ▲ | 10 | **2** | 3.1 | ***cysK*** | cysteine synthase A, O-acetylserine sulfhydrolase A subunit |
| WT | ▲ | 10 | **3** | 2.7 | ***glnH*** | glutamine transporter subunit |
| WT | ▲ | 10 | **4** | 2.7 | ***gatY*** | D-tagatose 1,6-bisphosphate aldolase 2, catalytic subunit |
| WT | ▲ | 10 | **5** | 2.5 | ***glnQ*** | glutamine transporter subunit |
| WT | ▲ | 10 | **6** | 2.5 | ***glnA*** | glutamine synthetase |
| WT | ▲ | 10 | **7** | 2.5 | ***glnP*** | glutamine transporter subunit |
| WT | ▲ | 10 | **8** | 2.3 | ***metN*** | DL-methionine transporter subunit |
| WT | ▲ | 10 | **9** | 2.2 | ***manZ*** | mannose-specific enzyme IID component of PTS |
| WT | ▲ | 10 | **10** | 2.1 | ***yjiY*** | putative transporter |
| WT | ▲ | 20 | **1** | 3.7 | ***pstC*** | phosphate transporter subunit |
| WT | ▲ | 20 | **2** | 3.3 | ***yjiY*** | putative transporter |
| WT | ▲ | 20 | **3** | 3.3 | ***alaC*** | valine-pyruvate aminotransferase 3 |
| WT | ▲ | 20 | **4** | 3.3 | ***gatY*** | D-tagatose 1,6-bisphosphate aldolase 2, catalytic subunit |
| WT | ▲ | 20 | **5** | 3.1 | ***uspA*** | universal stress global response regulator |
| WT | ▲ | 20 | **6** | 3.0 | ***gatA*** | galactitol-specific enzyme IIA component of PTS |
| WT | ▲ | 20 | **7** | 3.0 | ***clpB*** | protein disaggregation chaperone |
| WT | ▲ | 20 | **8** | 3.0 | ***pntA*** | pyridine nucleotide transhydrogenase, alpha subunit |
| WT | ▲ | 20 | **9** | 3.0 | ***ibpA*** | heat shock chaperone |
| WT | ▲ | 20 | **10** | 2.9 | ***pstS*** | periplasmic phosphate binding protein, high-affinity |

**Table S6. Host KEGG pathway enrichment results (CSV file).** KEGG pathway enrichment was identified using the DAVID tool. Enrichment was performed on datasets of significantly up- and downregulated host genes (FDR < 0.05) versus 0 min (= no phage). WT, wild type; hfq, Δ*hfq* mutant.

Column descriptions:

- STRAIN – *E. coli* host strain and time point (e.g., DE_P1_10_vs_nophage_hfq = differential expression upon P1vir infection at 10 min vs. no-phage control in the Δ*hfq* strain).
- Down/Up – direction of regulation of the gene set (downregulated or upregulated).
- Category – annotation category (here: KEGG pathway).
- TermID:Term – KEGG pathway identifier and pathway name combined in a single field.
- TermID – KEGG pathway identifier.
- Term – KEGG pathway name.
- Count – Number of genes from the input list mapped to the pathway.
- % – Percentage of input genes mapped to the pathway.
- PValue – Enrichment p-value for the pathway.
- Genes – List of genes from the input set associated with the pathway.
- List Total – Total number of genes in the input list.
- Pop Hits – Number of genes in the background population annotated to the pathway.
- Pop Total – Total number of genes in the background population.
- Fold Enrichment – Enrichment ratio of observed vs. expected hits in the pathway.
- Bonferroni – Bonferroni-adjusted p-value.
- Benjamini – Benjamini–Hochberg FDR-adjusted p-value.
- FDR – False discovery rate reported by DAVID.

**Table S7. Host GO term enrichment results (CSV file).** Gene Ontology (GO) enrichment across Biological Process (BP), Cellular Component (CC), and Molecular Function (MF) categories was identified usting the GSAn tool. Enrichment was performed on datasets of significantly up- and downregulated host genes (FDR < 0.05) versus 0 min (= no phage). WT, wild type; hfq, Δ*hfq* mutant.

Column descriptions:

- STRAIN – *E. coli* host strain and time point (e.g., DE_P1_10_vs_nophage_hfq = differential expression upon P1vir infection at 10 min vs. no-phage control in the Δ*hfq* strain).
- Down/Up – direction of regulation of the gene set (downregulated or upregulated).
- GOID – gene Ontology identifier of the enriched term.
- Name – name (label) of the enriched GO term.
- Ontology – GO category of the term (BP, CC, or MF).
- IC – Information content of the GO term (term specificity measure).
- Term_depth – depth of the GO term in the ontology hierarchy.
- Covered_genes – number of genes from the input set annotated to the GO term.
- Synthetic – indicator of whether the term is a synthetic/representative term generated by GSAn.
- Genes – list of genes from the input set associated with the GO term.

**Table S8. Hfq-favored motif scan of P1vir transcripts (CSV file).**

For each coding sequence (CDS), a 5′ window (−30 to +60 nt around the start codon) and a 3′ window (−40 to +20 nt around the stop codon) were scanned for Hfq-favored motif features. Gene-specific ΔΔlog2FC (Δ*hfq* vs. WT) at 10 and 20 min and induction ratio difference (Δ*hfq* vs. WT) at 10 and 20 min are also included.

Column descriptions:

- Gene symbol – gene name as given in the annotation (systematic name) and used throughout the text/figures.
- w5_len – length (nt) of the 5′ window scanned around the start codon (−30..+60 nt, oriented to the mRNA sense).
- w3_len – length (nt) of the 3′ window scanned around the stop codon (−40..+20 nt, oriented to the mRNA sense).
- AAN_repeats – number of AAN clusters (≥ 3 consecutive AAN triplets) in the 5’-window (distal-face motif; N = any nt).
- AAN_max_run – maximum number of consecutive AAN triplets in any cluster in the 5′ window.
- ARN_repeats – number of ARN clusters (≥ 3 consecutive ARN triplets) in the 5′-window (R = A/G).
- ARN_max_run – maximum number of consecutive ARN triplets in any cluster in the 5′ window.
- UA_repeats – number of (UA) tandem-repeat patches (≥ 3 repeats) in the 5′ window (lateral rim contact).
- UA_max_repeats – size of the longest (UA)_k patch in the 5′ window, reported as *k*.
- polyU_count – number of poly-U tracts (runs of U ≥ 5) in the 3′ window (proximal-face motif, often at intrinsic terminators).
- polyU_max – length of the longest poly-U tract in the 3′ window.
- ddlog2FC_10 – between-strain contrast at 10 min: ($\log_2 FC_{\Delta hfq} - \log_2 FC_{WT}$).
- ddlog2FC_20 – between-strain contrast at 20 min: ($\log_2 FC_{\Delta hfq} - \log_2 FC_{WT}$).
- IRD_10 – induction ratio difference (Δ*hfq* vs. WT) at 10 min, calculated as 2^(ddlog2FC_10).
- IRD_20 – induction ratio difference (Δ*hfq* vs. WT) at 20 min, calculated as 2^(ddlog2FC_20).

**Table S9. Statistics for candidate Hfq-sensitive motifs in P1vir transcripts.** For motif–effect analyses, we used the $\Delta\Delta\log_2FC$ across 10 and 20 min per gene and compared Motif-positive (+) versus Motif-negative (−) groups using the Mann–Whitney U test. Motifs used for statistical testing include:

- AAN_max_run – maximum number of consecutive AAN triplets in any cluster in the 5′ window;
- ARN_max_run – maximum number of consecutive ARN triplets in any cluster in the 5′ window;
- polyU_max – length of the longest poly-U tract in the 3′ window;
- UA_max_repeats – size of the longest (UA)_*k* patch in the 5′ window, reported as *k*;
- Any motif – presence of any of the above motifs.

| | Motif (+) | | | Motif (−) | | | Statistics | | | |
|---|---|---|---|---|---|---|---|---|---|---|
| **Motif** | ***N*** | **10 min** | **20 min** | ***N*** | **10 min** | **20 min** | **U (10 min)** | ***p* (10 min)** | **U (20 min)** | ***p* (20 min)** |
| **AAN_max_run** | 5 | −0.14 | −0.50 | 38 | −0.09 | −0.45 | 97 | 0.956 | 98 | 0.927 |
| **ARN_max_run** | 14 | −0.11 | −0.41 | 29 | −0.03 | −0.49 | 180 | 0.56 | 207 | 0.928 |
| **polyU_max** | 5 | −0.70 | −0.82 | 38 | −0.03 | −0.42 | 45 | 0.059 | 51 | 0.101 |
| **UA_max_repeats** | 1 | −0.09 | −0.42 | 42 | −0.11 | −0.49 | 21 | 1 | 23 | 0.930 |
| **Any motif** | 16 | −0.24 | −0.41 | 27 | −0.03 | −0.49 | 174 | 0.297 | 225 | 0.831 |
